# Mechanism and evolutionary origins of Alanine-tail C-degron recognition by E3 ligases Pirh2 and CRL2-KLHDC10

**DOI:** 10.1101/2023.05.03.539038

**Authors:** Pratik Rajendra Patil, A. Maxwell Burroughs, Mohit Misra, Federico Cerullo, Ivan Dikic, L. Aravind, Claudio A. P. Joazeiro

**Affiliations:** Center for Molecular Biology of Heidelberg University (ZMBH), DKFZ-ZMBH-Alliance, D-69120 Heidelberg, Germany; National Center for Biotechnology Information, National Library of Medicine, National Institutes of Health, Bethesda, MD 20894, USA; Institute of Biochemistry II, Goethe University Faculty of Medicine, Theodor-Stern-Kai 7, 60590, Frankfurt am Main, Germany; Buchmann Institute for Molecular Life Sciences, Goethe University Frankfurt, Max-von-Laue-Strasse 15, 60438, Frankfurt am Main, Germany; Department of Molecular Medicine, UF Scripps Biomedical Research, Jupiter, 33458, FL, USA

**Keywords:** Ala-tail, C-end degron, E3 ubiquitin ligase, KLHDC10, Pirh2, polyalanine, protein degradation, protein-protein interaction, protein quality control, structure prediction

## Abstract

In Ribosome-associated Quality Control (RQC), nascent-polypeptides produced by interrupted translation are modified with C-terminal polyalanine tails (‘Ala-tails’) that function outside ribosomes to induce ubiquitylation by Pirh2 or CRL2-KLHDC10 E3 ligases. Here we investigate the molecular basis of Ala-tail function using biochemical and *in silico* approaches. We show that Pirh2 and KLHDC10 directly bind to Ala-tails, and structural predictions identify candidate Ala-tail binding sites, which we experimentally validate. The degron-binding pockets and specific pocket residues implicated in Ala-tail recognition are conserved among Pirh2 and KLHDC10 homologs, suggesting that an important function of these ligases across eukaryotes is in targeting Ala-tailed substrates. Moreover, we establish that the two Ala-tail binding pockets have convergently evolved, either from an ancient module of bacterial provenance (Pirh2) or *via* tinkering of a widespread C-degron recognition element (KLHDC10). These results shed light on the recognition of a simple degron sequence and the evolution of Ala-tail proteolytic signaling.

## Introduction

Protein quality control is essential to maintain proteome homeostasis and cell viability.^1^ Central to this task, among other mechanisms, is the proteasomal degradation of aberrant proteins mediated through the action of E3 ubiquitin ligases, which confer specificity in ubiquitylation.^2, 3^ E3 ligases—of which there are over 650 in humans^4^—often target their specific substrates by recognizing linear motifs, or degrons, in the substrates. Degrons can be present internally, in the N-terminus, or in the C-terminus (C-end) of the substrate.^5^ C-end degrons are a newly discovered degron class, whose sequence diversity and cognate E3 ligases are only beginning to be elucidated. They can be naturally present in some proteins, be exposed as a result of proteolytic cleavage, be generated by damage, or be appended as a protein modification, such as in the protein quality control pathway known as Ribosome-associated Quality Control (RQC).^6–20^

RQC functions in eliminating incomplete polypeptides produced when ribosomes stall during protein synthesis.^21^ Dedicated factors sense and dissociate stalled ribosomes, only to generate large (60S) ribosomal subunits still blocked with nascent polypeptide-tRNA conjugates. In mammals, the protein NEMF recognizes such obstructed large subunits and recruits Ala-charged tRNA to extend the C-termini of the nascent polypeptides with a polyalanine tract (‘Ala-tail’).^8, 22, 23^ Ala-tailing can then mediate degradation of the nascent polypeptides through at least two routes. In the canonical RQC pathway (RQC-L), C-terminal elongation helps expose Lys residues otherwise buried in the ribosomal exit tunnel for on-ribosome ubiquitylation by the E3 ligase Listerin;^8, 24^ in the RQC-C pathway, Ala-tails themselves serve as C-end degrons that mediate interaction in an extra-ribosomal fashion with either Pirh2, an E3 ligase, or KLHDC10, the substrate recognition subunit of a Cullin-2 RING ligase (CRL2) complex.^8^

Pirh2 was originally characterized as a <u>p</u>53-induced <u>R</u>ING-<u>H2</u> domain containing E3 ligase (also known as Rchy1) that ubiquitylates p53 and promotes its proteolysis.^25, 26^ Among other known Pirh2 substrates are the tumor suppressor proteins p63 and p73, and the oncoprotein c-Myc.^27–29^ Exactly how Pirh2 recognizes these substrates is not understood. The biological roles of KLHDC10 also remain poorly understood.^30^ Mechanistically, in addition to targeting Ala-tailed substrates, KLHDC10 has been independently found to recognize C-end degrons ending in TrpGly, ProGly or AlaGly.^10^ The KLHDC10 paralog KLHDC2 recognizes degrons ending in di-Gly, and the paralog KLHDC3 binds to degrons ending in ArgGly, LysGly or GlnGly.^10^ Crystal structures of KLHDC2 bound to peptides corresponding to the C-end degron of prematurely terminated selenoproteins (SelK and SelS), and of a proteolytic fragment of the deubiquitinating enzyme USP1, have been reported,^12^ providing first insights into the structural basis for C-end degron recognition.

Here we study the mechanism of Ala-tail degron function. We show that the Ala-tail interacts directly with Pirh2 and KLHDC10 and use a combination of biochemical assays and *in silico* structural prediction to identify and experimentally validate unique Ala-tail binding sites in these E3 ligases. Importantly, the predicted interaction models provide insights into the basis for selectivity in Ala-tail degron sensing. Finally, we show that both the degron-binding sites and specific residues implicated in Ala-tail recognition are highly conserved among Pirh2 and KLHDC10 homologs that are widely distributed across the eukaryotic tree. Pirh2 can be confidently inferred as being present in the last eukaryotic common ancestor (LECA), suggesting that at least one E3 ligase targeting Ala-tail-containing substrates was functional since the origin of extant eukaryotes. We show that, whereas KLHDC10 evolved its substrate-binding pocket *via* subtle modifications to a comparable C-degron recognition site shared with its paralogs KLHDC2 and KLHDC3, the Pirh2 pocket shows a more dramatic history. It was derived from an ancient module of bacterial provenance that was reused *via* modification and augmentation to give rise to a unique binding interface.

## Results

### Reconstitution of Pirh2 and KLHDC10 binding to the Ala-tail degron in vitro

Pirh2 is comprised of three independently folding parts, namely the N-terminal module (NTM; amino acids 1-137), the RING domain (amino acids 138-189), and C-terminal domain (CTD; amino acids 190-261) (Figures 1A **and S1A**). Their structures in isolation have been solved using NMR spectroscopy (PDB accession codes 2K2C, 2JRJ and 2K2D, respectively).^26^ Our analysis showed that the NTM is a composite module comprised of three distinct types of Zn-chelating domains (**Figures S1B-E**): 1) the N-terminal-most binuclear Zn-binding CHY Treble-clef domain, distantly related to the ZZ and RING domains.^31^ 2) This is followed by a central classic Zn-ribbon (ZnR) domain^32^ which coordinates a single Zn ion. 3) The C-terminal part of the NTM comprises of three repeats of a Zn-binding domain, which again evolved from a standalone C-terminal fragment of a Treble-clef domain.^31^ Each of these repeats binds a single Zn ion in conjunction with one of the adjacent repeats. Thus, the entire NTM binds a total of 6 Zn ions.^26^ The CTD is centered on a single ZnR domain related to the central domain in the NTM and chelates a Zn ion. The NTM and the CTD together mediate p53 recognition,^26^ whereas the E3 ligase catalytic RING domain serves as the binding site for ubiquitin-carrying E2 conjugases.^4^

**Figure 1.**
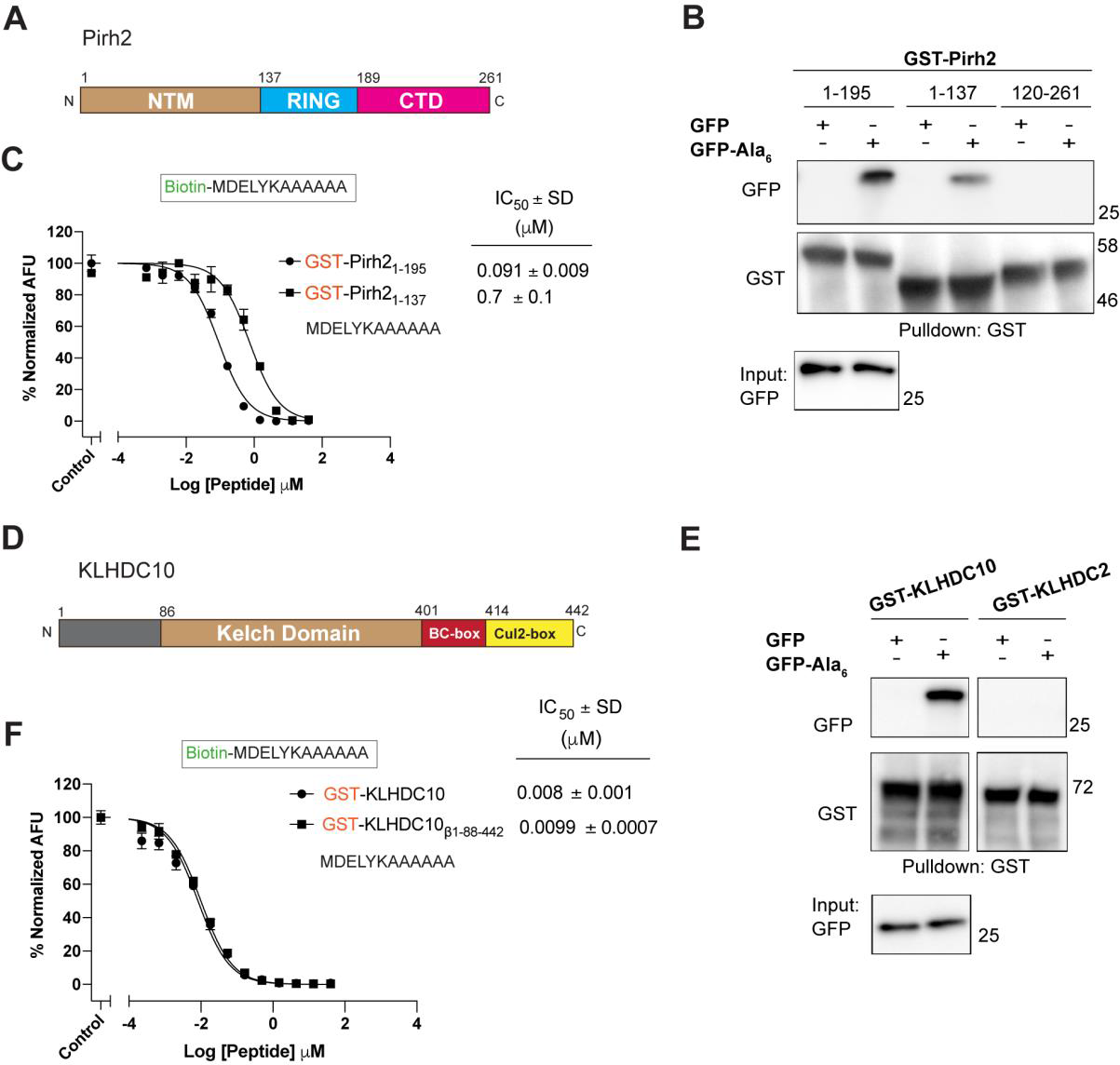
Pirh2 and KLHDC10 directly bind to Ala-tails. (A) Domain arrangement of human Pirh2: NTM, N-terminal Module; RING, ‘Really Interesting New Gene’ domain; CTD, C-terminal Domain. (B) *In vitro* GST pull-down assay using the specified GST-Pirh2 truncation constructs and recombinantly purified GFP or GFP-Ala_6_. Anti-GST and anti-GFP immunoblots are indicated. (C) AlphaScreen assay to assess Pirh2-Ala-tail binding. Reactants were GST fusions with Pirh2 fragments containing both the NTM and RING domain (Pirh2_1-195_) or the NTM alone (Pirh2_1-137_), and a biotinylated Ala-tail peptide (MDELYKAAAAAA). Proximity-induced fluorescence signal (see Methods) was monitored in presence of increasing amounts of a competing, non-biotinylated Ala-tail peptide to determine IC_50_ values from the dose response curve. Each data point is in triplicate, error bars represent ± SD. The *y*-axis presents the normalized AlphaScreen signal, as arbitrary fluorescence units (AFU). (D) Domain arrangement of human KLHDC10. (E) *In vitro* GST pull-down assay using the GST-KLHDC10 and GST-KLHDC2 constructs and recombinantly purified GFP or GFP-Ala_6_. Anti-GST and anti-GFP immunoblots are indicated. (F) AlphaScreen assay to assess KLHDC10-Ala-tail binding. As in panel C, but using GST-KLHDC10 and GST-KLHDC10_β1-88-442_ instead.

To begin mapping the parts of Pirh2 involved in Ala-tail recognition, we expressed and purified Pirh2 fragments fused to an N-terminal GST tag from *E. coli* and evaluated their binding to recombinant GFP-Ala_6_, a fusion protein which we had previously shown to be degraded *in vivo* in a Pirh2- and KLHDC10-dependent manner.^8^ The results of GST pulldown followed by anti-GFP immunoblot show that the Pirh2 NTM (Pirh2_1-137_) was sufficient for GFP-Ala_6_ binding (Figure 1B), although a construct containing both the NTM and RING domains (Pirh2_1-195_) pulled down GFP-Ala_6_ more efficiently than the NTM alone. On the other hand, a construct containing both the RING domain and CTD (Pirh2_120-261_) did not have detectable GFP-Ala_6_ binding activity. These results suggest that Pirh2 recognition of Ala tails is mediated by the NTM and is stimulated by the RING domain.

To both independently confirm and quantitate the interaction between the Ala-tail degron and Pirh2, we next developed an *in vitro* binding assay based on the Amplified Luminescence Proximity Homogeneous Assay (AlphaScreen). This assay relies on the proximity-dependent transfer of singlet oxygen from streptavidin-conjugated donor beads to anti-GST antibody-conjugated acceptor beads, which ultimately produces a fluorescence signal. To mediate bead proximity, we used GST-Pirh2_1-195_ for anti-GST bead binding and designed a biotinylated peptide for streptavidin bead binding. The peptide, MDELYKAAAAAA (herein referred to as ‘Ala-tail peptide’), comprises six Ala residues preceded by the last six amino acids of GFP, which include polar and charged residues to improve peptide solubility. We then validated the assay by examining fluorescence emission as indicative of Ala-tail peptide binding to GST-Pirh2_1-195_. Indeed, in presence of both interacting partners, a signal could be observed, which decreased upon titration of a competing, unlabeled (i.e., not biotinylated) Ala-tail peptide (Figure 1C). The dose-response curves provided a half maximal inhibitory concentration (IC_50_) value of ∼90 nM for the Ala-tail peptide-Pirh2_1-195_ interaction (Figure 1C). Finally, consistent with the pulldown experiments, deletion of the RING domain decreased Ala-tail binding by nearly 10-fold, with the Pirh2 construct containing the NTM alone (Pirh2_1-137_) having an IC_50_ of ∼700 nM for the Ala-tail peptide (Figure 1C).

We also examined Ala-tail binding to KLHDC10. This protein has a central Kelch β-propeller domain composed of six repeats of the Kelch motif (amino acids 86-401; Figures 1D **and S2**), flanked at the N-terminus by an 85-amino acid long region predicted to be mostly unstructured, and at the C-terminus, by a BC-box and a Cul2-box (amino acids 402-414 and 415-442, respectively), which together comprise a single trihelical domain.^33^ In the paralog KLHDC2, the Kelch domain recognizes substrate degrons,^12^ whereas the BC-box binds to the Cullin-2 adaptor proteins Elongin B (POZ domain) and Elongin C, and the Cul2-box provides an additional, direct interaction site for Cullin-2, which in turn recruits the RING domain protein Rbx1 into a CRL2 E3 ligase complex.^34, 35^ Similar interactions likely apply for KLHDC10,^33^ although we could not formally demonstrate that the Kelch domain suffices for C-degron binding, as proteins lacking the C-terminal BC-box and/or Cul2-box appeared to be unstable when expressed in *E. coli*. Nonetheless, we confirmed that recombinant KLHDC10, but not KLHDC2, specifically pulled down GFP-Ala_6_ (Figure 1E) and measured an IC_50_ of ∼10 nM for the Ala-tail peptide-KLHDC10 interaction in the AlphaScreen assay (Figure 1F).

### Evaluation of features of the Ala-tail degron implicated in E3 ligase binding

The above results provide strong evidence that Pirh2 and KLHDC10 bind directly and with high affinity to the Ala-tail degron. We next set out to identify features of the degron that confer specificity towards the E3 ligases. For this purpose, we performed titrations of Ala tail-variant, unlabeled peptides in the AlphaScreen assay to determine their IC_50_ values for GST-Pirh2 and -KLHDC10 fusion proteins.

For KLHDC10, we utilized a construct deleted for the unstructured N-terminal 87 amino acids, except for amino acids 40-51, which improved the protein’s expression and solubility (KLHDC10_β1-88-442_; **Figure S2**). Each Kelch repeat folds as a blade of the β-propeller, comprised of four β-strands that are designated A-D from the inner to the outer position (**Figure S2**); amino acids 40-51 were maintained in the KLHDC10_β1-88-442_ construct as they were predicted by AlphaFold to fold as β strand-D of the Kelch domain repeat 6. KLHDC10 full length and KLHDC10_β1-88-442_ proteins had similar IC_50_ values for Ala-tail peptide interaction in the AlphaScreen assay (Figure 1F).

In one set of experiments, we investigated the basis for our previous observation that polyAla sequences must be located at a protein’s very C-terminus to function as degrons.^8^ For example, extending an Ala5 sequence with a single Thr led to its complete loss of destabilizing activity *in vivo*, suggesting that polyAla-mediated protein degradation requires sensing of both a C-terminal Ala and its alpha-carboxyl group.^8^ Therefore, we used the AlphaScreen assay to test the effect of replacing the terminal alpha-carboxyl group of the Ala-tail peptide with an amide group. Strikingly, this substitution alone resulted in a two thousand-fold or greater reduction in the peptide affinity to both Pirh2 (Figure 2A) and KLHDC10 (Figure 2B), implying that these E3 ligases make critical contacts with the terminal alpha-carboxyl group of the degron.

**Figure 2.**
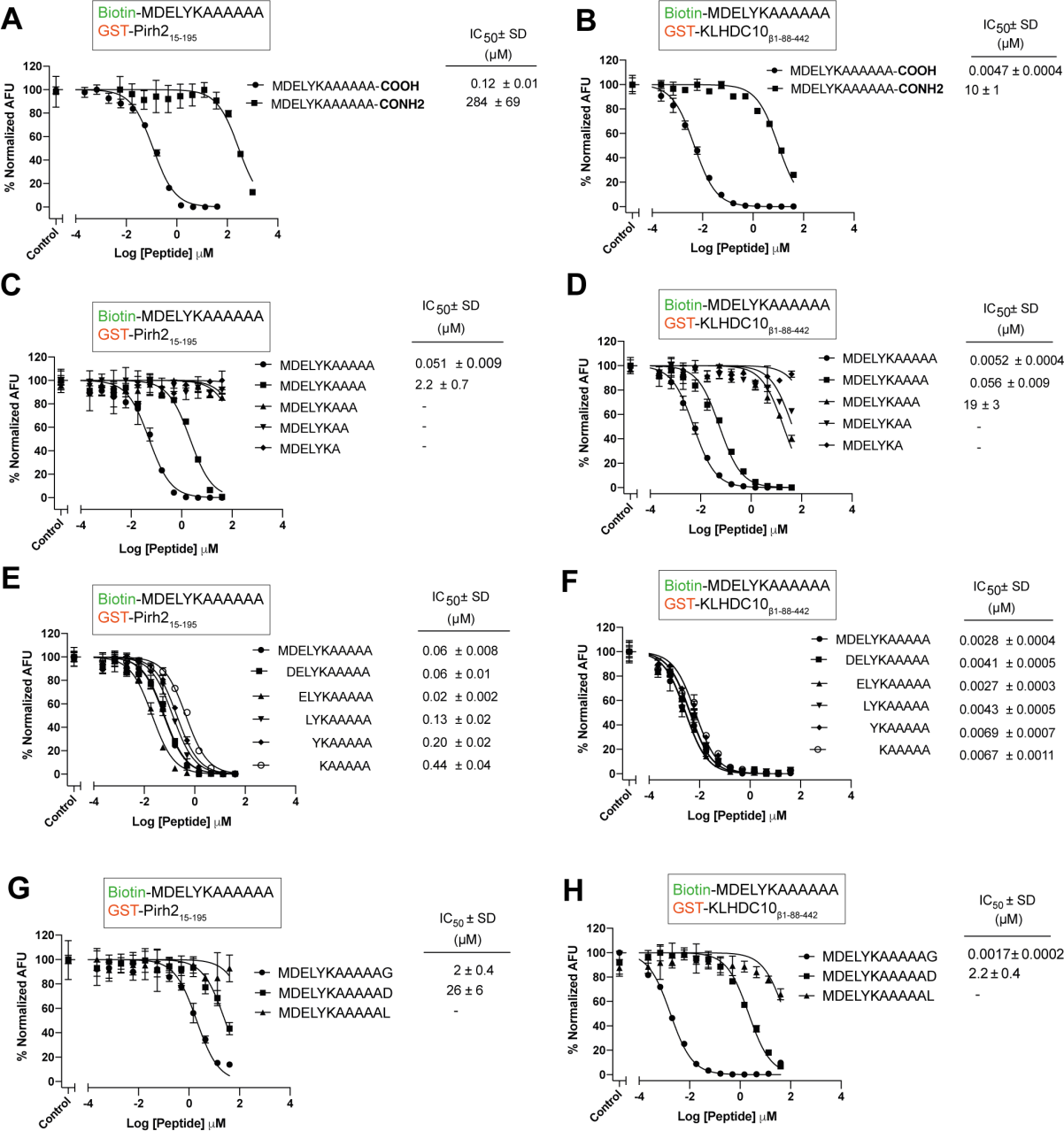
Four Ala residues at the C-terminus and the terminal carboxyl group are essential for Ala-tail degron binding by Pirh2 and KLHDC10. (A-H) As in Figure 1C, but using GST-Pirh2_15-195_ (A, C, E, G), GST-KLHDC10_β1-88-442_ (B, D, F, H) and competing peptides, as indicated, to measure the effects of: C-terminal amide substitution (A, B); 1-5 Ala residues at the C-terminus (C, D); shortened N-terminal sequences in Ala_5_-containing peptides (E, F); 5 Ala followed by a C-terminal Gly, Asp or Leu (G, H).

In another set of experiments, we investigated the basis for the observation that at least four Ala residues are required for a homopolymeric Ala-tail to act as a degron *in vivo*^8^ by determining the number of Ala residues in a degron peptide required for binding to each of the E3 ligases. The results show that a peptide containing five C-terminal Ala had an IC_50_ of ∼50 nM for Pirh2 binding, while the affinity decreased to ∼2 µM when only four Ala were present (Figure 2C). Further shortening the tail caused binding to become undetectable under the assay conditions. Similar observations were made for KLHDC10, in that Ala-tail peptides containing five or four Ala residues at the C-terminus bound strongly (IC_50_ of 5 nM and 56 nM, respectively) whereas a peptide with three Ala showed a further 400-fold reduction in binding (IC_50_ of 19 μM; Figure 2D). This loss of binding was not due to the overall peptide length, as deleting residues from the N-terminus of the Ala-tail peptide representing the last six amino acids of GFP only affected binding to Pirh2 (Figure 2E) and KLHDC10 (Figure 2F) to a much lesser extent.

Finally, we investigated the requirement for the residue in the most C-terminal position (defined as −1), which is critical for recognition by C-end E3 ligases. For example, KLHDC2 binds to C-terminal Gly degrons, TRIM7 favors Gln-end degrons, and FEM1 binds Arg-end degrons.^12, 14, 17, 18^ The effect of replacing Ala −1 in the Ala-tail peptide with Gly, Leu or Asp was examined. Gly has a smaller side chain than Ala and lacks hydrophobicity. Leu, on the other hand, has a hydrophobic side chain but which is larger than that of Ala. Lastly, Asp has a small but negatively charged side chain. The results in Figure 2G show that, in contrast to the Ala-tail peptide that binds to Pirh2 with an IC_50_ in the 50-120 nM range, the IC_50_ for a peptide with the same sequence except for ending in Gly was ∼2 μM, whereas Asp at −1 position further decreased the IC_50_ to ∼26 μM and the presence of Leu at −1 completely abolished binding under the assay conditions. These results suggest a strict requirement for Ala at −1 position for an Ala-tail degron to be recognized by Pirh2.

In contrast to Pirh2, in addition to substrates containing homopolymeric Ala tails,^8^ KLHDC10 had been previously found to bind to substrates ending in -TrpGly, -ProGly and -AlaGly.^9, 10, 12^ Consistent with this, using the AlphaScreen assay we determined that KLHDC10 binds to an Ala-tail peptide variant ending in Gly −1 with high affinity (IC_50_ ∼2 nM; Figure 2H), which was comparable to KLHDC10’s affinity for the ‘parental’ peptide ending in Ala. However, and as was the case for Pirh2, Ala-tail peptides ending in Asp or Leu bound to KLHDC10 with markedly reduced affinity (nearly five hundred-fold lower and to an undetectable level under the assay conditions, respectively).

In summary, the above results identify features of the Ala-tail degron that confer specificity towards Pirh2 and KLHDC10. We show that the polyAla sequence must be located at a protein’s very C-terminus to function as a degron, that a minimal of four Ala residues at the C-terminus as well as the terminal carboxyl group are both essential for Ala-tail binding, and that Ala at −1 position is required for binding to Pirh2 while, in line with previous reports by others,^9, 10, 12^ KLHDC10 can also accept a peptide with Gly at −1.

### AlphaFold2 predicts Ala-tail degron binding sites in Pirh2 and KLHDC10

We next sought to understand the structural basis for Ala-tail degron recognition by Pirh2 and KLHDC10. *In silico* analyses utilizing AlphaFold2^36, 37^ proved insightful in predicting the structure of complexes of Pirh2 or KLHDC10 with the Ala-tail peptide and delineating general molecular principles underlying the interactions (Figures 3 and 4).

**Figure 3.**
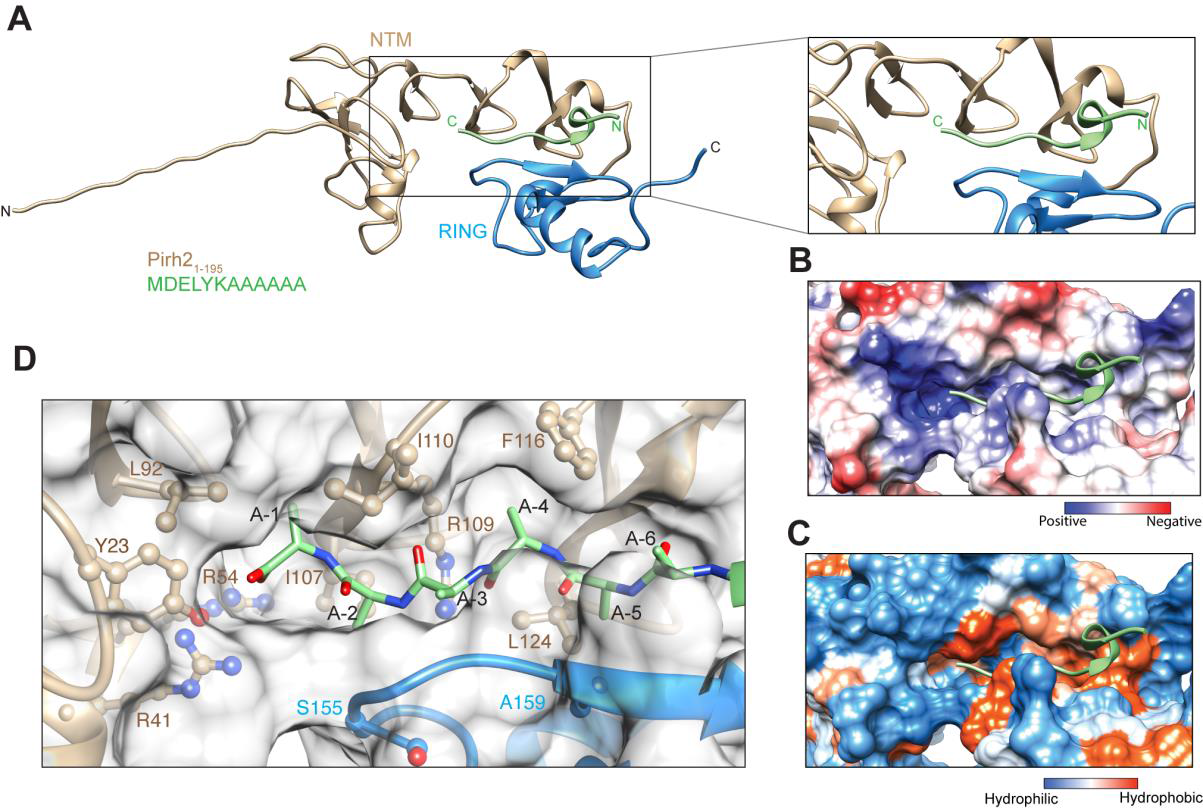
Identification of a candidate Ala-tail C-degron binding site in Pirh2. (A) AlphaFold2 prediction of Pirh2_1-195_ in complex with an Ala-tail peptide. The highest-ranking among predicted models is shown. Ribbon representation of Pirh2 NTM and RING domains colored according to Figure 1. The Ala-tail peptide is shown in green. The inset shows the Ala-tail degron binding site in close-up view. (B, C) Electrostatic surface potential and (C) surface hydrophobicity represented for the focused region shown in A; Ala-tail peptide in green. (D) A close-view of the Ala-tail peptide in complex with Pirh2_1-195_, with the C-terminal Ala residues displayed in stick representation and colored in green. Pirh2_1-195_ is represented as a transparent surface, with residues in vicinity of the peptide labelled and sides chains in ball-stick representation.

**Figure 4.**
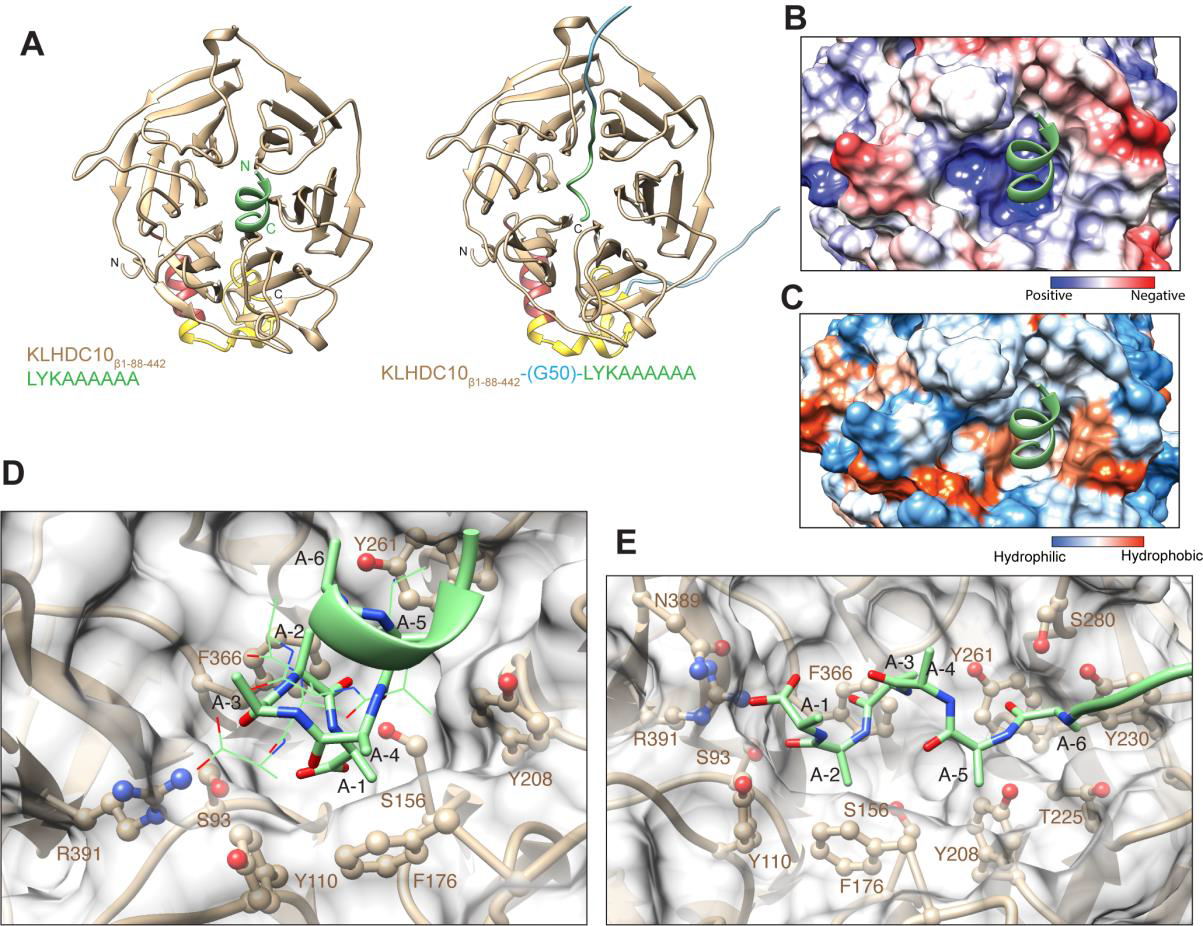
Identification of a candidate Ala-tail C-degron binding site in KLHDC10. (A) As in Figure 3, but for KLHDC10_β1-88-442_. KLHDC10 colored as in Figure 1D. (A) Top predictions of KLHDC10_β1-88-442_ in complex with a LYKAAAAAA peptide (*left*) or fused to the LYKAAAAAA peptide through a Gly50 linker (*right*). (B, C) Electrostatic surface potential and (C) surface hydrophobicity represented for the focused region shown in A *left*; Ala-tail peptide in green. (D) Close-up view of the *left* panel shown in ‘A’. Two orientations of Ala-tail peptide (green) are presented, in stick and wire representations. (E) Close-up view of the *right* panel shown in A.

In one analysis, Pirh2_1-195_ and Ala-tail peptide (MDEYLKAAAAAA) sequences were input as separate chains. In another, an Ala-tail was connected to the C-terminus of Pirh2_1-195_ *via* a long and flexible Gly_50_ linker. In both analyses, the 5 top-ranking models generated by AlphaFold2 predicted almost identical Ala_6_ peptide binding modes (**Figures S3A and S3B**). The highest-scoring models out of the top-ranking predictions obtained from the experiment with Pirh2 and the degron peptide sequence provided as separate chains are discussed further.

Superposition of the previously reported NMR structures of Pirh2’s individual NTM and RING domains^26^ with the AlphaFold2 model of Pirh2_1-195_ revealed an RMSD of ∼1.8 Å, validating the accuracy of the computed model (**Figure S3C**). Close inspection of the Cys and His residues that form the NTM zinc finger motifs as well as the RING domain confirmed that all are oriented similarly in the AlphaFold2 model and the NMR structures (**Figure S3C**), which is remarkable given the absence of coordinated zinc ions in the former.

In the AlphaFold2-predicted structure of full-length Pirh2, the NTM and RING domains establish close contacts (**Figure S1A**). The interaction is mediated by a hydrophobic interface between NTM residues Leu124, Leu126, Leu130 and Ile137 and RING domain Val140, Val161 and Leu167. In addition, several residues contribute polar interactions. Arg109 and Lys135 from the NTM hydrogen bond with main chain carbonyl of His153 and Cys164 of the RING domain, respectively; Ser141, in the linker between the NTM and RING domain, hydrogen bonds with the NTM Asn123 main chain carbonyl and the RING domain His153; and the NTM His22 main-chain carbonyl is poised to make a hydrogen bond with the RING domain Arg156 side chain. Strikingly, in the predicted structure of the Pirh2-peptide complex, the Ala-tail peptide binding site lies in a pocket formed by the NTM at the interface with the RING domain (Figure 3A). This model is consistent with our biochemical analyses that uncovered the NTM domain as being both required and sufficient for binding, and with the RING domain playing a stimulatory role in the process (Figures 1B and 1C).

The predicted Ala-tail binding site in Pirh2 is a narrow channel where the Ala-tail peptide fits precisely, in an extended, flexible loop-like conformation. The C-terminus of the peptide reaches deep into the binding pocket, while the N-terminus lies on the shallow open end. The binding pocket has features that are suitable for Ala-tail binding—it is amphipathic, with hydrophilic and hydrophobic residues dispersed throughout, and has a basic patch which complements the C-terminal alpha-carboxyl group of the Ala-tail peptide as well as the main chain carbonyl groups of the peptide backbone. (Figures 3B and 3C). The C-terminus of the Ala-tail peptide lies in the vicinity of Pirh2’s polar Tyr23 and positively charged Arg41and Arg54 (Figure 3D), whose side chains are suitably positioned to establish hydrogen bonds and/or electrostatic interactions with the degron’s obligatory, negatively charged terminal alpha-carboxyl group. The side chain of Ala −1 of the degron peptide fits into a shallow hydrophobic cleft formed by Leu92 and Ile110 of the NTM. The space constraint at the cleft, together with its hydrophobic nature, adequately supports Pirh2’s preference for the small, hydrophobic side chain of Ala at position-1. The side chains of Ala −2, −4 and −5 are likewise all enclosed into small clefts placed throughout the binding pocket, lined by Arg109 and hydrophobic residues Ile107, Phe116 and Leu124. Ala −3 is placed such that its side chain points away from the binding pocket towards relatively open space, suggesting that this position of degron peptide may be more amenable for substitution. Consistently, we had previously reported that the *B. subtilis* SsrA tag (which ends in ALAA) could be bound by Pirh2 *in vitro*^38^ although with moderate affinity (IC_50_ ∼470 nM; **Figure S4**). Lastly, Ala −6 also orients its side chain away from the pocket. Thus, the binding pocket in Pirh2 perfectly accommodates the last five Ala of the degron peptide, which is consistent with the strongest binding by the Ala-tail peptide containing five Ala in the AlphaScreen assay (Figure 2C).

Our biochemical analyses showed that the RING domain was not essential for, but stimulated Ala tail binding (Figures 1B and 1C). The structure suggests mechanisms underlying the cooperativity effect. First, the RING domain contributes a lateral surface to the Ala-tail binding pocket, thereby helping exclude water to create a more hydrophobic environment. Moreover, the Ala-tail peptide backbone runs parallel to and establishes contacts with a beta sheet of RING domain involving several main chain interactions, such as the hydrogen bonding between the main chain carbonyl group of Ser155 and the amide of Ala-2, and between both backbone carbonyl and amide groups of Ala159 and the Ala-5 main chain.

For KLHDC10_β1-88-442_, AlphaFold2 predicted the Ala-tail degron binding site at the ‘top’ surface of the Kelch domain (defined as the side of a beta-propeller containing the loops linking beta-strands A to D, and B to C; ^39^). However, in the initial analyses the resulting peptide orientations were not uniform (data not shown) and in the highest ranked model, the alpha-carboxyl group of the degron did not make any contacts with KLHDC10. Therefore, we ran new predictions using an Ala-tail peptide with fewer GFP-derived N-terminal residues to prevent potential ‘distraction’ caused by those sequences (**Figure S5**). With this strategy, the predicted binding mode in all complexes, regardless of the approach used, consisted of the Ala-tail peptide bound on the top surface, with the C-terminal Ala penetrating deep in the central pocket (Figures 4A, **S5A and S5B**). Remarkably, co-crystallization studies had revealed an equivalent site in the Kelch domain of KLHDC2 for binding to the di-Gly C-end degron.^12^ Notably, in the model of apo-full length KLHDC10, the N-terminus of the protein, whose sequence has two Ala, at positions 3 and 4, is predicted to fold back and also bind to the same pocket as the Ala-tail peptide. This interaction is reminiscent of the recently reported C-degron mimicry by the C-terminus of KLHDC2, which results in autoinhibition.^20^ Whether the KLHDC10 interaction is real and is Ala-dependent, and whether it provides a regulatory mechanism for substrate binding has yet to be investigated.

Although all analyses pointed to the same mode of Ala-tail binding to KLHDC10, specific features of the computed models were more heterogeneous than for Pirh2. In the top prediction, KLHDC10 recognized the Ala-tail degron in a coiled conformation, stabilized through intra-chain hydrogen bonds between backbone carbonyl and amide groups (Figure 4A, ***left***). This prediction is consistent with the helical nature of polyAla sequences ^40, 41^ but a helical structure has also been observed for other C-end degrons with unrelated sequence when bound to their cognate E3 ligases.^12, 14, 15, 18^ Due to this alpha-helical conformation, the Ala side chains of the degron point towards the wall of the binding site in a spiral arrangement, with those of Ala-1 and Ala-2 flanked against the narrowed hydrophobic surface of the binding pocket. In contrast, the top scoring prediction when the Ala-tail peptide was fused to KLHDC10 through a Gly_50_ linker, the Ala tail-degron bound KLHDC10 in a non-helical, disordered conformation (Figure 4A, ***right***).

In either case the central pocket in KLHDC10 appears favorable for Ala-tail recognition, as a basic patch is found at its deep bottom and hydrophobic residues line its wall (Figures 4B and 4C). The C-terminal carboxyl group of the degron sits in the vicinity of KLHDC10 Ser93, Ser156 and Arg391 residues (Figure 4D), suitably placed to form hydrogen bonds and/or electrostatic interactions; in two out of five predictions, Tyr110 is in proximity to the C-terminal carboxyl group, Tyr208 hydrogen bonds with the backbone amide of Ala-5, and Tyr261 hydrogen bonds with the main chain carbonyl of Ala-6 of the Ala-tail peptide. The binding pocket at this end is narrowed by the bulky, hydrophobic side chains of KLHDC10 Phe176 and Phe366, which flank Ala −1 and −2 of the degron peptide, respectively. Steric hindrance caused by such bulky residues is a plausible explanation for why Ala- or Gly-ending degrons are recognized by KLHDC10. The pocket is relatively wide at the top, such that there are minimal contacts beyond the Ala-tail peptide Ala −4, which is flanked by KLHDC10 Phe176. On the other hand, in a model where Ala-tail peptide binds in a non-helical conformation, the high affinity for the Ala-tail degron is supported by the selectivity towards Ala residues. The Ala-tail peptide Ala-2 is flanked by Phe176, Ala-3 with Phe366, Ala-5 with Tyr208 and Ala-6 with Thr225 and Tyr230 (Figure 4E). The terminal carboxyl in this model is stabilized by Arg391, Asn389 and Tyr110. Additionally, Tyr208 and Ser280 form hydrogen bond with the main chain amide of Ala-5 and carbonyl of Ala-6, respectively.

Therefore, using AlphaFold2 we have been able to identify candidate Ala-tail binding pockets in both Pirh2 and KLHDC10 that are overall consistent with critical biochemical features implicated in degron binding from the perspectives of both the E3 ligases and the degron.

### Validation of the Ala-tail degron binding sites in Pirh2 and KLHDC10 by analysis of rationally designed mutations

We next tested the AlphaFold2 predicted Ala-tail binding sites in Pirh2 and KLHDC10 by introducing single amino acid mutations in the E3 ligases and examining their effect on Ala-tail binding using the GST pulldown assay.

The results in Figure 5A show that Pirh2 mutations affecting polar, charged or hydrophobic residues on the peptide binding pocket either abolished (Y23A, R41A, R109A, L124D) or decreased (R54A) GFP-Ala_6_ pulldown. As described above, Tyr23, Arg41 and Arg54 are located deep in the binding pocket and are predicted to interact with the carboxyl terminus of the Ala-tail degron; the Arg109 side chain forms a hydrogen bond with the backbone carbonyl group of Ala −4; and Leu124 flanks Ala −5. The specificity of the results is further highlighted by the finding that one of the mutants we designed (I107D; Figure 5A) as well as mutants affecting several residues that are not part of the predicted binding pocket (not shown) did not have an obvious defect in GFP-Ala_6_ binding under our assay conditions.

**Figure 5.**
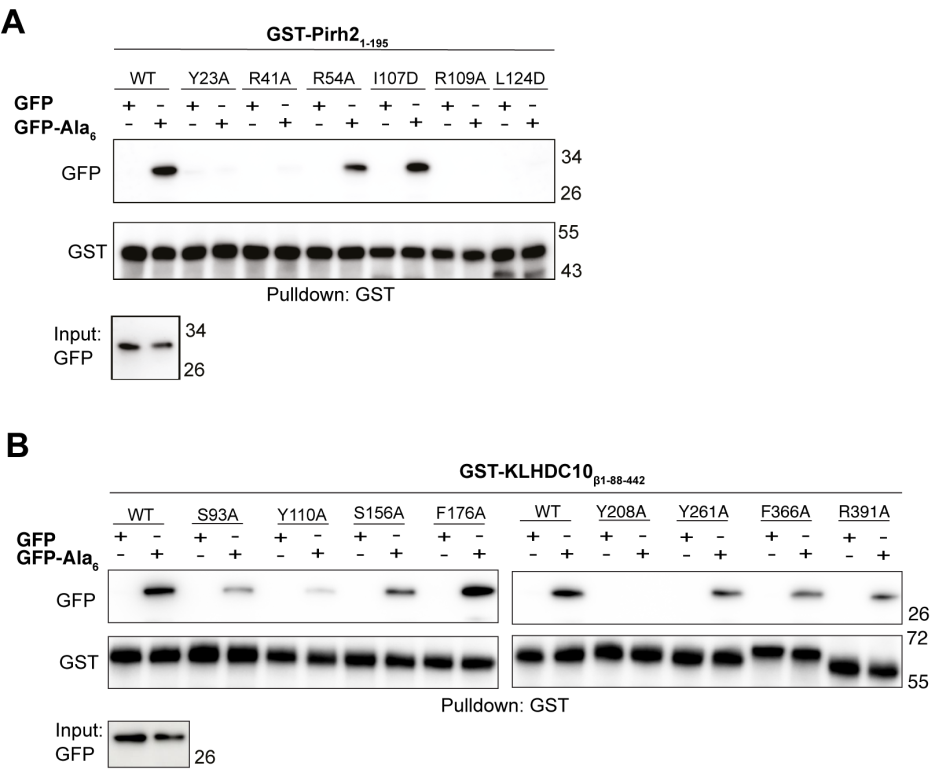
Validation of residues implicated in Ala-tail degron recognition by Pirh2 and KLHDC10. (A) *In vitro* GST pull-down assay using GST-Pirh2_1-195_ WT and mutant constructs as indicated, and recombinantly purified GFP or GFP-Ala_6_. Anti-GST and anti-GFP immunoblots are shown. (B) As in A, but for GST-KLHDC10_β1-88-442_ WT and indicated mutant constructs.

Also for KLHDC10, several mutations had a negative effect on the ability to pull down GFP-Ala_6_ (Figure 5B). That was the case for mutations of Ser93, Tyr110, Ser156, and Arg391 to Ala, all expected to interfere with interaction with the degron’s C-terminal carboxyl group; Phe366, which may interact with either Ala-2 or Ala-3 of the degron; Tyr261, which may interact with Ala-6; and Tyr208, predicted to interact with the main-chain amide of Ala-5, whose mutation had the strongest detrimental effect observed.

Therefore, the results of rational mutagenesis experiments provide evidence in support of the computationally predicted Ala-tail binding sites in Pirh2 and KLHDC10.

### Conservation of the C-degron binding site in Pirh2 and KLHDC10 homologs

We next examined whether the C-degron binding sites identified in Pirh2 and KLHDC10 are shared among their homologs.

A multiple sequence alignment (MSA) shows that several Pirh2 residues predicted to create the degron binding pocket and to bind the Ala-tail peptide are indeed highly conserved across all major eukaryotic lineages with available sequence data (Figure 6A). Of the inferred Ala-tail binding residues, the positions corresponding to Tyr23, Arg41 and Arg54 are all derived from the N-terminal-most CHY Treble-clef domain of the NTM (**Figure S1**). The first of these residues is almost always an aromatic residue (Tyr or Trp) in all Pirh2 homologs and is structurally localized adjacent to the residues corresponding to Arg41 and Arg54, which are both retained as positively charged positions in most Pirh2 homologs. This is consistent with the notion that the aromatic position engages in cation-π interactions with the two positively charged residues which stabilizes the end of the binding pocket and provides a chemical “counter-part” for the terminal carboxyl group of the Ala tail. Mapping the degree of conservation of surface residues (which include residues implicated both in binding to the alpha-carboxyl group of the Ala-tail and in creating an amphipathic pocket) on the human Pirh2 structure revealed a clear clustering of residue conservation at the deep end of the Ala-tail binding pocket (Figure 6B). Underscoring the relevance of this observation, the clustering of highly conserved Pirh2 residues was also observed for the RING domain surface implicated in binding to E2 conjugases as part of the E3 ligase catalytic mechanism (Figure 6B). Finally, the identified Ala-tail binding pocket was structurally conserved at the interface of the NTM and the RING domain in AlphaFold2 predictions of all Pirh2 homologs examined from fission yeast to humans (Figure 6C). Together, these observations suggest that Pirh2 homologs in other organisms are equipped for functioning in Ala-tail binding. In support of this possibility, we have found evidence that the most distant and divergent sequence analyzed in Figure 6A, that of the *S. pombe* Pirh2 homolog, selectively interacts with Ala-tails, as its GST fusion pulled down GFP-Ala_6_ but not wild type GFP or GFP fused to C-terminal Thr_6_ or (Ala-Thr)_3_ tails (Figure 6D).

**Figure 6.**
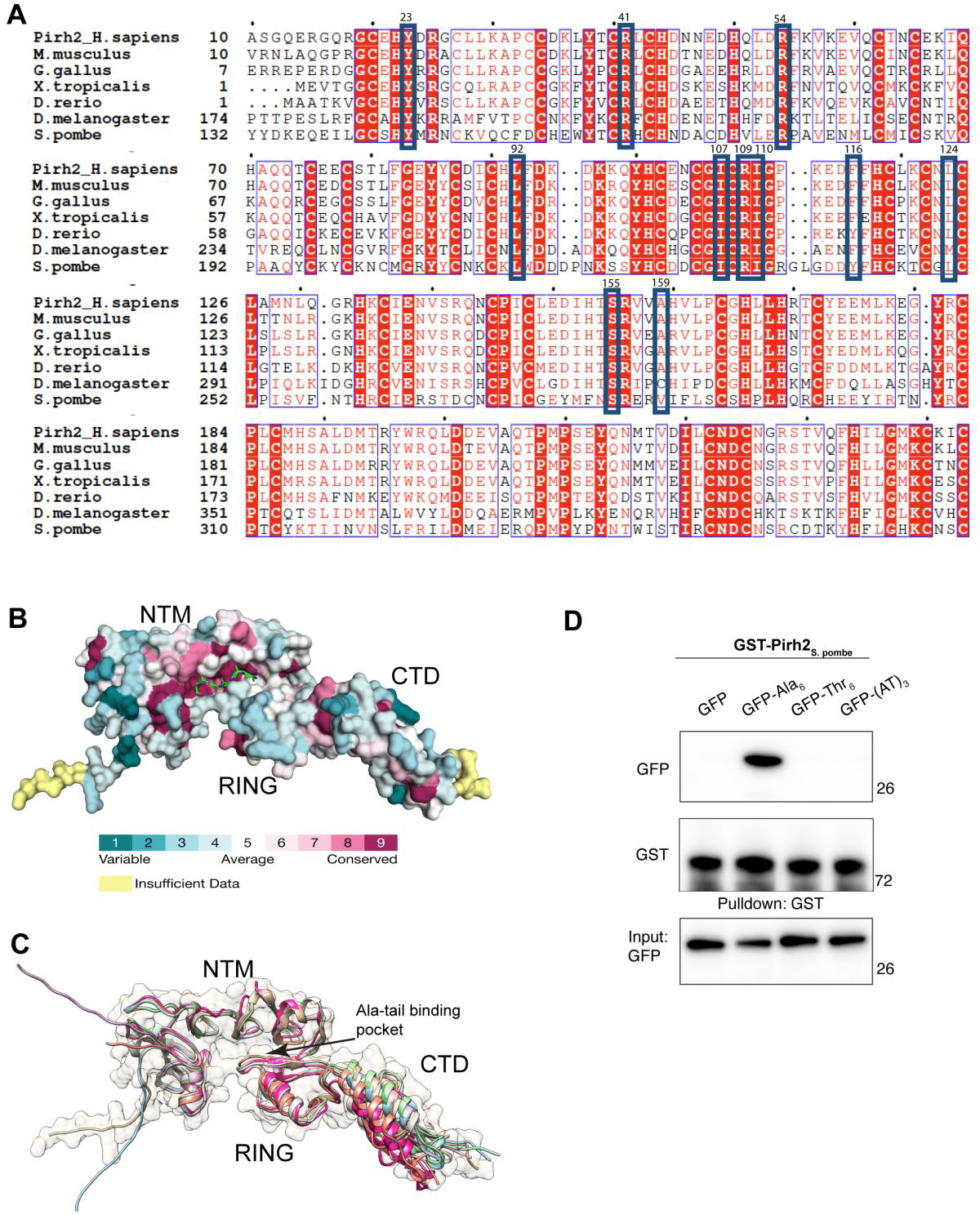
Conservation of residues implicated in Ala-tail binding among Pirh2 homologs. (A) Multiple sequence alignment for Pirh2 homologs. The blue boxes highlight residues predicted to stabilize Ala-tail peptide at the binding pocket in Pirh2 that are shown in Figure 3D and corresponding residues in the homologs. (B) Residues conserved in Pirh2 homologs mapped onto the AlphaFold structure of human Pirh2 (UniProt: Q96PM5), using ConSurf webserver. The color key for variable to conserved residues is shown below. Peptide in stick representation at the Ala-tail binding site. (C) AlphaFold models for Pirh2 homologs in (A) are superposed. Backbone of each protein is in ribbon representation in different colors, for human Pirh2 surface is also represented with transparency. (D) *In vitro* GST pull-down assay using GST tagged full length Pirh2 homolog from *S. pombe* and recombinantly purified GFP, GFP-Ala_6,_ GFP-Thr_6_ or GFP-(Ala-Thr)_3_. Anti-GST and anti-GFP immunoblots are indicated.

To analyze the evolution of KLHDC10 we generated a phylogenetic tree of homologs, which allowed us to precisely delineate KLHDC10 from KLHDC2 and KLHDC3, the defining members of paralogous sister clades (Figures 7A**, S6 and S7**). KLHDC10 has a more restricted distribution than Pirh2, being present across animals and their sister groups, namely choanoflagellates, filastereans and ichthyosporeans. They are also present in some other eukaryotic lineages like stramenopiles and alveolates but appear to have been lost in fungi. Analysis of the KLHDC10 orthologs reveals conservation of both sequence (Figure 7B) and the predicted Ala-tail-binding pocket (Figure 7C), and includes the residues implicated in the degron binding (e.g., Ser93, Ser156, Phe176, Tyr208, Thr225, Tyr261, Phe366 and Arg391).

**Figure 7.**
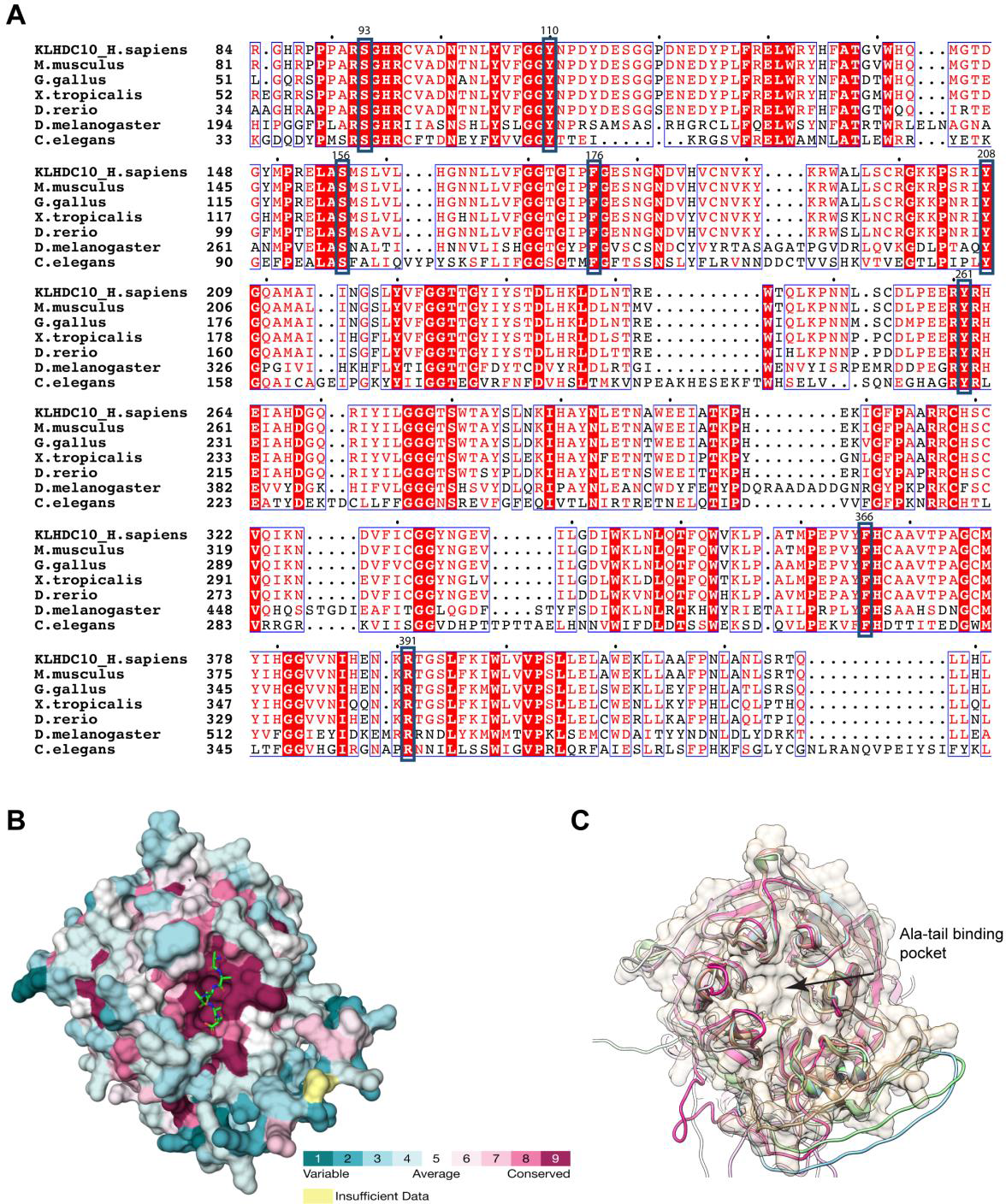
Conservation of residues implicated in Ala-tail binding among KLHDC10 homologs. (A-C) As in Figure 6, but for the KLHDC10 Kelch domain. In (A) blue boxes highlight residues implicated in Ala-tail binding shown in Figure 4D.

Finally, and consistent with the distinct degron-binding specificities of other human KLHDC10-family proteins that are most closely related to KLHDC10, such as KLHDC2, the residues predicted to mediate Ala-tail contacting sites in the binding pocket are strongly conserved in the KLHDC10 clade suggesting that the Ala-tail binding specificity is likely to be retained across most members of the clade. Three of these positions, namely those corresponding to Ser93, Tyr110 and Phe366 (as an aromatic position), are also conserved in KLHDC2, a sister clade of KLHDC10 (**Figures S6, S7 and S8**). Hence, we propose that these residues might play a more general role in recognizing C-degrons across the clades, whereas those positions conserved only in KLHDC10 are likely to mediate the specific recognition of Ala-tails generated in the RQC-C pathway (Figures 7 and S8).

**Figure 8.**
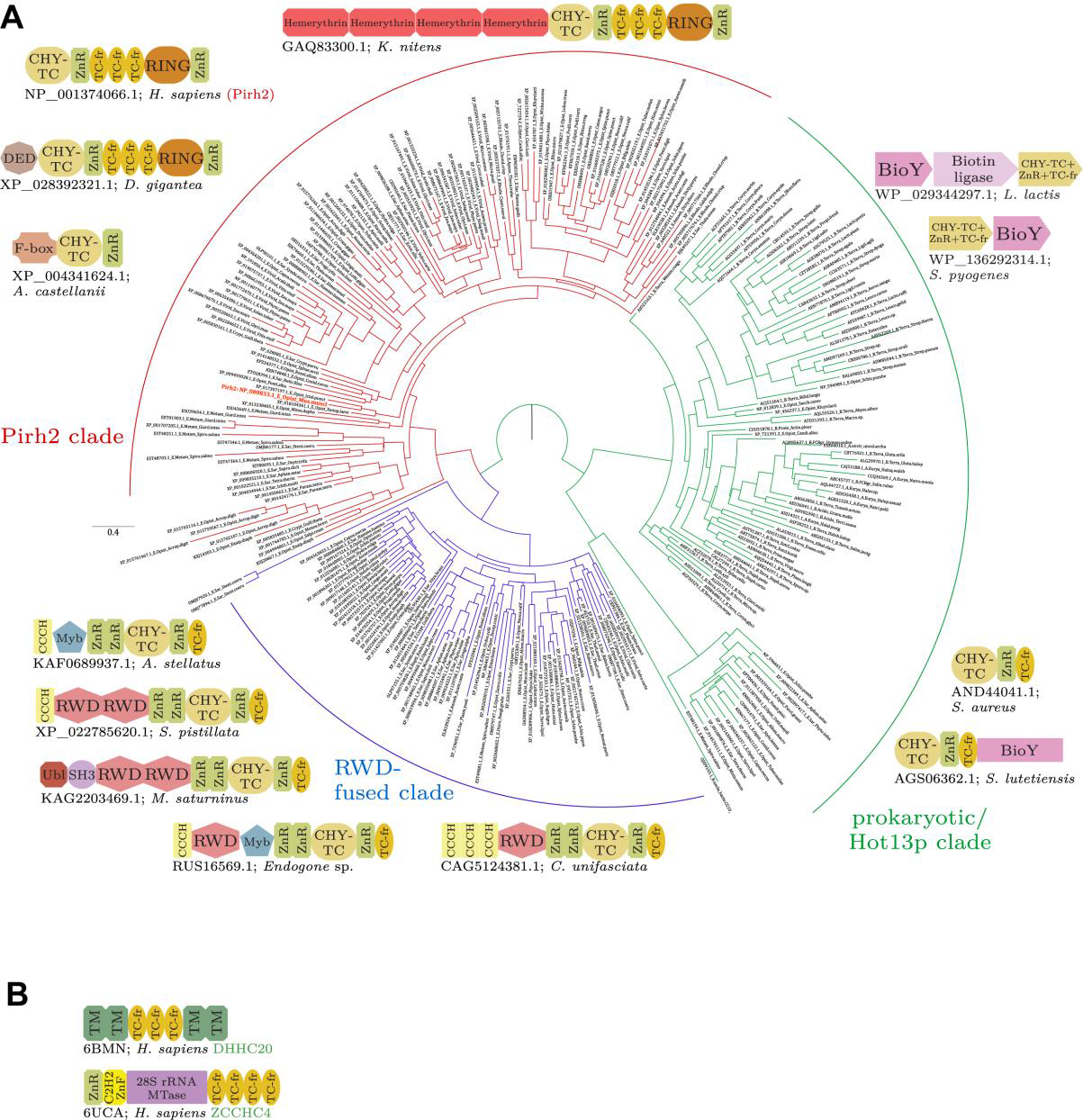
Phylogenetic distribution of domains in Pirh2 NTM and examples of human Treble-clef fragment repeat-containing proteins. (A) Phylogenetic tree of the CHY-TC (Treble-clef) and ZnR module found in the NTM. Monophyletic clades are colored and labeled. Representative domain architectures and gene neighborhoods are provided next to their corresponding clades in the tree. The mouse Pirh2 sequence is highlighted in bold lettering and colored red. (B) Domain architectures of human Treble-clef fragment repeat-containing proteins.

The above results provide strong support for the predicted binding sites for the Ala-tail degron in Pirh2 and KLHDC10. Moreover, the results implicate Pirh2 and KLHDC10 homologs in having a critical, conserved function in sensing Ala-tails and related C-end degrons.

### Evolutionary affinities and origins of the Ala-tail recognition modules of Pirh2 and KLHDC10

Pirh2 and KLHDC10 are unrelated proteins, indicating that Ala-tail binding activity emerged at least twice in eukaryotic evolution. Our analyses indicate that Pirh2 orthologs are widely distributed across most major eukaryotic lineages. While there is some uncertainty about the eukaryotic tree of life, the presence of Pirh2 in the “greater plant” lineage (Archeplastida) and lineages such as Heterolobosea (e.g., *Naegleria*) and Metamonada (*Giardia* and *Spironucleus*), which are considered basal branches in the eukaryotic tree, makes it likely that Pirh2 was present in the last eukaryotic common ancestor (LECA) (Figure 8). This ancestral version can be reconstructed as already having the tripartite organization with the composite NTM, the central RING and C-terminal ZnR. Moreover, it is predicted as conserving most of the key Ala-tail recognition residues (**Figure S9**).

In contrast, we found that KLHDC10 has a patchier distribution, being present primarily in animals, their immediate sister groups and outside of them in some lineages of the Stramenopile-Alveolate-Rhizarian clade (SAR) (**Figure S7**). This makes it less likely of being inherited from the LECA. Its paralog, the KLHDC2 clade, also shows a limited phyletic distribution being present primarily in animals and Amoebozoans. However, our analyses indicate that the KLHDC3 clade is more widespread, with a presence in animals and their closest sister groups, amoebozoans, SAR and plants. Thus, KLHDC3 has greater likelihood of being a more ancient clade of Kelch-repeat C-degron binding subunits (but cannot be confidently inferred as being in the LECA), with KLHDC10 and KLHDC2 being derived later as part of the radiation of versions that acquired distinct specificities. Hence, it might be postulated that the more widespread Pirh2 is probably the original Ala-tail-specific E3 component inherited from the LECA.

However, the presence of two analogous E3 ligases with overlapping specificities might have allowed the loss of either version in certain lineages. For example, our analyses suggest that Pirh2 has been secondarily lost in certain animals such as *C. elegans* and kinetoplastids. At least in *C. elegans*, the presence of a member of the KLHDC10 clade likely allows the use of Ala-tail dependent C-degrons (Figure 7). In contrast, *Saccharomyces cerevisiae* lacks both Pirh2 and KLHDC10, which might relate to the presence of Ala-Thr-tails instead of pure Ala-tails in this organism.^23^ A similar situation is seen across Basidiomycete fungi, which have lost both Pirh2 and KLHDC10. It is also notable that several eukaryotic clades display unusual modifications of the ancestral Pirh2 system (Figure 8): 1) In amoebozoans, the tripartite NTM is combined with a F-box instead of a RING domain, suggesting that the same Ala-tail recognition module is now routed for action *via* an E3 ligase with an F-box subunit. 2) Across the “greater-plant” clade the Pirh2 proteins are fused to 4 copies of the heme-binding hemerythrin domain, suggesting that the RQC-C pathway might interface with sensing of light or redox stimuli *via* these domains. 3) In the basal animal lineage of the cnidarians, Pirh2 is fused to N-terminal Death-effector domains (DED) that mediate homotypic interactions specific to apoptosis in animals. Hence, it is possible that in this case the same pathway is specifically recruited for degrading proteins as part of the programmed cell death process.

These observations raise the question of the origin of the unique tripartite NTM of the Pirh2 proteins in the first place. Our sequence profile searches recover significant similarity to not just other Pirh2 proteins but also a range of other proteins from eukaryotes, bacteria and archaea (Figure 8). The common denominator across these proteins is the N-terminal CHY Treble-clef and the central ZnR domain (together defined as the CHY-finger module in Pfam) and possibly including a rudimentary version of a single repeat of the Treble-clef fragment (Figures 8 and S9). A minimal version comprised of just this core element occurs as a standalone protein in several bacteria, certain archaea (Haloarchaea) and was secondarily transferred to fungi (e.g., *S. cerevisiae* Hot13). There is no evidence in these organisms for the coupling of this core version of the NTM with RQC or prokaryotic ubiquitin-like-conjugating systems. However, we found both conserved operonic linkages and direct fusions in the same polypeptide to the biotin transporter BioY and the biotin ligase (BirA) suggesting that it might function as an adaptor that aids in biotinylation or regulating biotin transport/conjugation by binding the unusual hydrophobic cytoplasmic tail of the BioY transporter. In contrast, the yeast version Hot13 has been shown to specifically bind the protein Mia40 the receptor for import of proteins into the mitochondrial intermembrane space and sequester Zn ions from it.^42^

Our analysis showed that the minimal version of the NTM was acquired from bacteria and incorporated into two distinct ubiquitin-related functions by the time of the LECA (Figures 8 **and S9**). In one case it was fused to a RING domain resulting in Pirh2 and in the other, to the RWD domain and nucleic-acid-binding CCCH domains (and sometimes additionally DNA-binding MYB domains) (Figure 8). The RWD domain of the latter proteins might mediate interactions with E2 domains or specific nucleoprotein complexes, including ribosomes, while the NTM cognate is expected to be involved in peptide recognition, as in the case of Pirh2. A comparison of the inferred binding site in the core of the NTM between Pirh2, the Hot13-like clade and the RWD-fused clade shows that whereas the aromatic position equivalent to Tyr23 is conserved across all of them, the two key basic positions, equivalent to Arg41 and Arg54 are not. They are both aromatic positions in the Hot13 like clade, suggesting a distinct binding site in the form of an aromatic cage. One of the basic residues is replaced by a conserved acidic amino acid in the RWD-fused clade. Thus, the Ala-tail binding site appears to have emerged specifically in Pirh2 from a distinct ancestral binding site inherited from the bacterial versions. Only in Pirh2, the peptide-binding site of the NTM was further augmented by triplication of the C-terminal Treble-clef fragment (**Figures S1B-E**). This triplicated element additionally acquired an independent existence in eukaryotes by being coupled to transmembrane (TM) segments, giving rise to the DHHC domain palmitoyltransferase catalytic domain,^43^ and as the C-terminal RNA-binding domain of the 28S rRNA methyltransferase ZCCHC4 (**Figures S1E and** 8B). The former version, in addition to binding the palmitoyl coA substrate, binds the linker peptide connecting it to the TM-segment in a manner comparable to the binding of the Ala-tail by Pirh2 (Figure 8).

## Discussion

We had previously reported that Ala-tails synthesized at the C-termini of ribosomal stalling products by the mammalian RQC pathway act as C-degrons that are targeted by the E3 ligases Pirh2 and CRL2-KLHDC10.^8^ Here, we analyze the E3 ligase-Ala tail interactions biochemically and generate structural predictions of their complexes using AlphaFold2. Together, the results provide insights into how a simple, homopolymeric polyAla degron sequence is selectively recognized by E3 ligases and shed new light onto the evolution of Ala tails as a proteolytic signal.

### AlphaFold as a prediction tool for protein-peptide complexes

AlphaFold has been successful in predicting protein structures solely based on protein sequences^36^ and its application has more recently been expanded towards predicting co-structures of protein-peptide complexes.^44, 45^ Indeed, using AlphaFold2, here we have generated reliable models for the complexes of Pirh2 and KLHDC10 with Ala-tail peptides, with several lines of evidence supporting the predictions: First, the predicted binding modes provide plausible explanations for experimentally determined features of both the E3 ligases and the degron that are implicated in the interactions (Figures 1 and 2). Second, site-specific mutagenesis to test the contribution of residues predicted to mediate degron binding support the overall structural models (Figures 3, 4, **and** 5). Third, the degron-binding pockets identified for both E3 ligases, relevant chemical features on the surfaces of those pockets, and specific residues predicted to mediate Ala-tail binding, are strongly conserved in evolution (Figures 6 and 7).

We note that, although the proposed role of individual E3 ligase residues implicated in binding has been substantiated through mutagenesis and evolutionary conservation analyses, our analyses are unable to distinguish among two highly ranked models describing specific KLHDC10 complexes, showing different conformations of the bound degron. Moreover, also for Pirh2 it is conceivable that the computational predictions provide an incomplete picture of the atomic interactions—for example, water molecules absent in the AlphaFold predictions might contribute to Ala-tail binding to Pirh2 and KLHDC10. Therefore, future experimental structure determination will be aimed at generating new models and verifying details of the proposed binding mode.

### The predicted binding modes provide plausible explanations for experimentally determined features of both the E3 ligases and the degron that characterize the interactions

Both Pirh2 and KLHDC10 bind to the Ala-tail degron with high affinity. Such strong interaction is comparable to that of several other E3 ligase-degron pairs, such as for KLHDC2 binding to its cognate C-end degrons.^12^ Zheng and colleagues have proposed that high affinity binding could have evolved to degrade substrates of low abundance, to eliminate substrates quickly, and/or to keep substrates at a low level due to their potential toxicity;^12^ such proposed roles of high-affinity substrate binding would also be relevant for the function of Pirh2 and KLHDC10 in protein quality control.^8^ Consistent with the experimentally determined binding affinities, the AlphaFold2 models for the Pirh2 and KLHDC10 complexes with the Ala tail degron are mediated by well-defined pockets with extensive interfaces. Moreover, the Ala-tail degron C-terminus is fully concealed from all sides in the deep binding pocket of KLHDC10, while in Pirh2 the degron C-terminus is not entirely buried in the pocket, providing a plausible explanation for the stronger Ala-tail binding by KLHDC10 compared to Pirh2.

The structural predictions also provide a plausible molecular explanation for the selective Ala-tail recognition by Pirh2 and KLHDC10 based on the degron’s sequence-, length- and COOH terminus-dependence for function. The binding pockets can fit 5 residues from the degron and establish continuous contacts with all those residues, consistent with our observation that high-affinity binding of both E3 ligases to Ala-tails requires at least four C-terminal Ala, and that binding affinity increases further with the presence of 5 Ala in the degron. For Pirh2, the side chains of Ala −1, −2, −4 and −5 are all enclosed into small clefts. Overall, the surfaces of the binding pockets in both E3 ligases contain several aliphatic and aromatic residues which complement the hydrophobicity of the degron’s Ala side chains. Notably, the choice of Ala −1 of the degron is specified by a hydrophobic cleft at both Pirh2 and KLHDC10 binding sites and is constrained by the small size of those clefts, that cannot accommodate larger side chains. Similar selectivity mechanisms to accommodate the terminal degron residue have been reported for other E3 ligases. For example, FEM1 stabilizes the terminal Arg in its cognate C-degron by an acidic cleft at the binding site.^15^ The structures also suggest that the free extreme C-terminal backbone COOH group plays a role in binding to both Pirh2 and KLHDC10, in agreement with the requirement for a polyAla tract to be present at the substrate’s very C-terminus to act as degron (^8^ and Figures 2A and 2B). The C-terminal COOH group is aligned in surrounded by polar and charged residues in the binding pockets of both E3 ligases. Similar distribution of polar residues can be observed in the experimentally determined crystal structures of other E3 ligases in complex with their cognate C-end degron peptides.^12, 14, 17, 18^ Overall, the electrostatic surface of both Pirh2 and KLHDC10 at the polyAla binding pocket appears highly electropositive (Figures 3B and 4B). consistent with the prediction that both the C-terminal carboxyl group and the protruding backbone carbonyl groups of the Ala-tail peptide are coordinated by either positively charged side chains or backbone amides.

Finally, the structural model of the Pirh2 complex predicts close intramolecular contacts between the Pirh2 NTM and RING domain, and that the Ala-tail peptide binding pocket lies in the NTM at the interface with the RING domain (Figure 3A). The model thus consistent with the stimulatory effect of the Pirh2 RING domain in Ala-tail binding to the NTM, as observed in our biochemical analyses (Figures 1B and 1C).

### Recognition of non-RQC substrates by Pirh2 and KLHDC10

Apart from Ala-tailed proteins modified by RQC activity, Pirh2 can artificially bind to the *B. subtilis* SsrA degron,^38^ which ends in with an Ala-tail-like sequence (ALAA). The human proteome lacks native proteins with more than three consecutive alanine residues at a cytosolic-exposed C-terminus, making it difficult to predict native full-length proteins that are likely to be targeted by Pirh2 or KLHDC10 *via* the mechanism described here. However, GFP fusion to the C-terminal 23 residues of TRAPPC11, which ends in 3 Ala (-MDDTSIAAA), sufficed to render the reporter unstable *in vivo* (in contrast with GFP-Ala3, which was only partially unstable under the same conditions) and expression of the fusion protein was restored to normal levels upon depletion of Pirh2 and KLHDC10.^8^ These results suggest that, in addition to targeting homopolymeric Ala tails, these E3 ligases may use the same mechanism to target native proteins harboring Ala-rich C-end degrons. Besides those, it is conceivable that Ala tail-like degrons might also result from specific or aberrant proteolytic cleavage.

Several substrates have been independently reported for both Pirh2^25–29^ and KLHDC10,^33^ none of which has a C-terminal sequence reminiscent of an Ala-tail. Biochemical analyses suggest that the tumor suppressor p53 primarily binds to CTD of Pirh2, with the NTM providing transient interaction.^26^ Likewise, Pirh2’s NTM and CTD work together in binding to c-Myc.^26, 29^ The last residue in c-Myc is an Ala but the corresponding peptide (KHKLEQLRNSCA) did not show detectable binding towards both Pirh2 or KLHDC10 in the AlphaScreen assay (**Figure S10**). Given that the CTD contains a ZnR domain that is conserved across all Pirh2 proteins (Figures 8 **and S1D**), it is conceivable that it constitutes a distinct binding side either by itself or in conjunction with the NTM to recognize a separate set of substrates from the C-terminal degrons.

### Evolutionary implications of Ala-tail signaling *via* Pirh2 and KLHDC10

Ala-tailing has a deep evolutionary history, but the mechanisms by which it activates protein degradation widely differ across the branches of the tree of life. NEMF/RqcH homologs, which mediate C-terminal tail synthesis, are tRNA- and ribosomal 60S subunit-binding proteins represented across all domains of life.^46^ Accordingly, bacteria have a RqcH-mediated Ala-tailing mechanism^46^ that is related to the yeast Rqc2-mediated Ala/Thr- and mammalian NEMF-mediated Ala-tailing reactions.^8, 47, 48^ Thus, the Ala-tailing modification mediated by an ancestor of NEMF/RqcH/Rqc2 proteins is presumed to have already existed in the Last Universal Common Ancestor (LUCA).

In the common ancestor of all bacteria, an additional mechanism for tagging the products of stalled ribosome for proteolysis appeared – the tmRNA/SsrA pathway. Although SsrA and RQC pathways are mechanistically unrelated, both modify aberrant nascent chains produced by incomplete translation with a C-terminal degron: polyAla in bacterial RQC,^46^ or an 8–35 residue-long peptide ending in ALAA in most bacteria, the SsrA tag.^49^ Moreover, these mechanisms have convergently evolved such that both tags are directly recognized by the ClpXP protease.^46, 49^

In eukaryotes, with the expansion of the ubiquitin system, further convergent ubiquitin-dependent mechanisms arose for targeting stalled polypeptides extended with C-terminal tails for degradation. One of these mechanisms involved use of the tail for exposing Lys residues buried in the exit tunnel for ubiquitylation *via* the Ltn1 ligase system on the 60S subunit (the RQC-L branch of the pathway; ^8^). As Ltn1 is found across most major eukaryotic lineages, C-terminal tailing-dependent Ltn1 activity is likely to have been already present in the LECA.

The results presented here show that at least one E3 ligase of the RQC-C branch of eukaryotic RQC pathway, *viz*., Pirh2, was also likely present in the LECA. In this branch, Ala-tailing has an extra-ribosomal degron function analogous to what is seen in bacteria.^8^ This in turn predicts that the NEMF orthologs across eukaryotes have an Ala-tailing function going in parallel with the presence of E3 ligases that specifically recognize homopolymeric Ala tails. Keeping with this, as noted above, the loss of both Ala-tail binding ligases in *S. cerevisiae,* likely relates to the random incorporation of both Ala and Thr, in the Rqc2-dependent tailing reaction.^23^ In addition to exposing Lys residues to support Ltn1,^23, 24, 50^ C-terminal tails containing both Ala and Thr are more prone to forming amyloid aggregates compared to Ala alone,^22^ raising the possibility that *S. cerevisiae* has evolved a further variant RQC activity that mediates protein aggregation to either sequester aberrant translation products, or enable their elimination *via* aggrephagy.^51^

Finally, our analysis unravels the evolutionary trajectories for the convergent origin of two distinct Ala tail-recognition modules in eukaryotic E3s. The less-widespread KLHDC10 evolved a binding pocket by subtle evolutionary tinkering of a preexisting C-degron recognition pocket that it partly shares with its paralogs KLHDC2 and KLHDC3. In contrast, Pirh2 represents a more dramatic case, wherein a composite module comprised of three distinct domains of ultimately bacterial provenance, was acquired prior to the LECA, and repurposed as an Ala tail-recognition module. This process involved both subtle modification of a preexisting ancient binding pocket, which is also utilized in other functional contexts, as well as its augmentation *via* triplication of the Treble-clef fragment domain.

In conclusion, the results provide further evidence for a widely conserved role of Ala-tails as a tag for proteolysis and pave the way for future biochemical and functional analyses of a wider range of organisms regarding these binding modules and protein modification mechanisms.

## Acknowledgments

We thank S. Pfeffer, R. Wade, M. Mayer, S. Filbeck, S. Richter and the Joazeiro lab for helpful discussions and comments on the data. We thank X. Zhao for assistance with KLHDC10 purification and A. Thrun, T. Dallinger and M. Bobeldijk for constructs. This project was supported in part by R01 grant NS102414 from the National Institute of Neurological Disorders and Stroke (NINDS) of the NIH (to C.J.; the content is solely the responsibility of the authors and does not necessarily represent the official views of the National Institutes of Health) and EU/EFPIA/OICR/McGill/KTH/Diamond Innovative Medicines Initiative 2 Joint Undertaking (EUbOPEN grant no 875510) (to I.D.). L.A. and A.M.B. are supported by intramural funds of the National Library of Medicine of the NIH.

## Author contributions

P.R.P.: Investigation; formal analysis; methodology; writing-original draft; writing-review and editing. A.M.B.: Investigation; formal analysis; writing-review and editing. M.M.: Methodology, formal analysis, writing-review and editing. F.C.: Investigation; writing-review and editing. I.D.: Supervision; writing-review and editing. L.A.: Formal analysis; supervision; writing-review and editing. C.A.P.J.: Conceptualization; resource; supervision; writing-original draft; writing-review and editing.

## STAR METHODS

### Resource availability

#### Lead contact

Claudio A. Joazeiro

#### Materials availability

All reagents generated in this study are available from the lead contact.

#### Data and code availability

This paper does not report original code. Any additional information required to reanalyze the data reported in this paper is available from the lead contact upon request.

#### Experimental model and subject details

##### Bacterial strain for recombinant protein expression

E. coli BL21 (DE3) cells were used to express recombinant proteins. Cells were transformed with respective plasmids and grown in LB media with required antibiotics as explained below.

#### Method details

##### Plasmid constructs

As described previously for Pirh2 and KLHDC10,^8^ KLHDC2 cDNA was subcloned into a modified pET15b vector that carries an N-terminal GST tag sequence. Truncation constructs of Pirh2 and KLHDC10 were generated by amplifying each fragment by PCR and subcloned into pET15b vector with a N-terminal GST tag sequence. The gene sequence coding for the full-length *S. pombe* Pirh2 homolog was synthesized (IDT) and subcloned into the pET15b vector with a N-terminal GST tag. KLHDC10_β1-88-442_ was generated by deleting the flexible region at the N-terminus (amino acids 1-39) and the internal loop (amino acids 52-87) of KLHDC10. KLHDC10_β1-88-442_ corresponds to the first β strand (amino acids 40-51) connected by a linker (GSGSG) to the rest of the protein (amino acids 88-442). The codon-optimized sequence with flanking NdeI and XhoI restriction sites was generated by IDT and subcloned in the pET15b vector with an N-terminal GST tag. The codon-optimized, bicistronic insert encoding Elongin B (amino acids 1-104) and Elongin C (amino acids 1-112) cloned in pET28b was described previously.^8^ For Pirh2 and KLHDC10 mutants, single point mutations were introduced into GST-Pirh2_1-195_ and GST-KLHDC10_β1-88-44_ in pET15b by site-directed mutagenesis. All plasmid constructs were verified by sequencing.

##### Recombinant protein purification

For expression of Pirh2 constructs, *E. coli* BL21 (DE3) cells were grown in LB media in presence of ampicillin (100 μg/ml), 10 mM MgCl_2_ and 10 μM ZnCl_2_. For expression of KLHDC10 and KLHDC2 constructs, *E. coli* BL21 (DE3) cells were co-transformed with pET15b vector carrying KLHDC10 and KLHDC2 and pET28b vector carrying a bicistronic insert for the expression of Elongin B (1-104) and Elongin C (1-112) as described before.^8^ Cells were grown in LB media with ampicillin (100 μg/ml) and kanamycin (50 μg/ml). For expression of all recombinant proteins, cells were grown until OD 0.8 at 37 °C, followed by overnight incubation at 18 °C in presence of 0.75 mM IPTG. On the following day, cells were harvested and stored at −80 °C until further processing.

GST-tagged Pirh2 proteins were purified by cell lysis using sonication in ice-cold lysis buffer (50 mM Tris pH 7.5 at 4 °C, 150 mM NaCl, 1 mM DTT, 10 µM ZnCl_2_, 1% Triton X 100 and EDTA-free protease inhibitor (Roche)). Lysates were clarified by centrifugation and supernatants were incubated with glutathione agarose bead slurry (Machery-Nagel) prewashed in wash buffer (50 mM Tris pH 7.5 at 4 °C, 150 mM NaCl, 1 mM DTT, 10 µM ZnCl_2_). After 2 h incubation at 4 °C, beads were washed three times using wash buffer. Bound GST-Pirh2 was eluted from beads by glutathione-containing elution buffer (50 mM Tris pH 7.5 at 4 °C, 150 mM NaCl, 1 mM DTT, 10 µM ZnCl_2_, 10 mM reduced glutathione). To remove glutathione, protein fractions were pooled and buffer-exchanged with wash buffer using Amicon Ultra-4, 10 kDa filters. Concentrated protein was aliquoted and stored at −80 °C.

For purification of GST-tagged KLHDC10 and GST-KLHDC2 proteins, cells were lysed by sonication in ice-cold lysis buffer (50 mM Tris pH 7.5 at 4 °C, 150 mM NaCl, 1 mM DTT, 0.5% Triton X 100 and EDTA-free protease inhibitors (Roche)). Proteins were purified using glutathione agarose bead as described above except that buffers did not contain ZnCl_2_. Buffer-exchanged, concentrated protein fractions were aliquoted and stored at −80 °C. BL21 cells co-transformed with KLHDC10_β1-88-442_ and Elongin B/C constructs were grown in TB media at 37 °C until O.D. 1, when the culture was cooled down to 18 °C and protein expression induced with 0.4 mM IPTG. After 18 h, cells were harvested and re-suspended into 50 mM Tris-HCl pH-7.5, 300 mM NaCl, 10% glycerol, 1 mM PMSF, 0.1 % lysozyme, 5 mM DTT and Roche protease inhibitor tablets. Cells were lysed by passing twice through a microfluidizer under 1,500 bar, at 4 °C. The lysate was centrifuged for 1 h at 4 °C at 15,000 x *g*. The supernatant was passed through a 0.8 µm filter and incubated with glutathione resin pre-equilibrated with re-suspension buffer. Beads were washed with the same buffer except for using a higher salt concentration (500 mM NaCl). The protein was eluted with 50 mM Tris-HCl pH 7.5, 500 mM NaCl, 15 mM glutathione and 5 mM DTT. Fractions containing the proteins of interest were pooled and diluted to 50 mM NaCl concentration using 50 mM Tris-HCl pH 7.5. This sample was subjected to anion exchange chromatography (Hiprep Q column) using a gradient between buffer A (50 mM Tris-HCl pH 7.5, 50 mM NaCl, 5 mM DTT) and buffer B (50 mM Tris-HCl pH 7.5, 1 M NaCl, 5 mM DTT). The peak fractions containing the proteins of interest were pooled and concentrated to 1 ml using Centricon filter (10 kDa cut-off). This sample was then used for size exclusion chromatography using Superdex 16/600 75pg. The complex of GST-KLHDC10 and Elongins B-C was concentrated to 1 mg/ml and flash frozen at −80 °C for AlphaScreen assays used to evaluate Ala-tail peptides.

Following a previously reported protocol, GFP, GFP-Ala_6_, GFP-Thr_6_ and GFP-(Ala-Thr)_3_ were purified from *B. subtilis* Δ*clpP* strains grown in LB media at 37 °C until OD 2.0.^46^ Cells were harvested and lysed as above in 50 mM Tris pH 8.0 at 4 °C, 150 mM NaCl, 0.5% NP40, 1 mM DTT. Clarified lysates were mixed with GFP-Trap agarose slurry by rotating for 1 h at 4 °C. Beads were washed in buffer without DTT. Elution was performed using 0.2 M glycine pH 2.5 at 4 °C, and the pH was neutralized by adding 1M Tris base pH 10.4. Aliquots of purified protein were stored at −80 °C with 10% (v/v) glycerol.

##### *In vitro* binding assay

Binding assays were performed using recombinantly purified proteins GST-Pirh2 or GST-KLHDC10 and GFP-Ala_6_.^8^ Briefly, glutathione agarose beads (Machery-Nagel) were washed and blocked with 1% BSA in assay buffer (20 mM Tris pH 7.5, 100 mM NaCl, 0.1% Tween-20) for 2 h at 4 °C. Glutathione beads were conjugated with 0.5-10 μg GST-Pirh2 or GST-KLHDC10 by incubating 10 μl BSA-blocked beads together with the respective protein in 100 μl assay buffer at 4 ^0^C. Following 2 h of incubation, beads were washed twice in assay buffer to remove excess unbound protein. Protein-conjugated beads were then incubated with 0.25 μg of either GFP or GFP-Ala_6_ in 100 μl assay buffer supplemented with 0.1% BSA for 1 h at 4 °C. Beads were then washed thoroughly to remove unbound protein and were resuspended in sample buffer. Binding of GFP-Ala_6_ was detected by anti-GFP and anti-GST blots.

##### Immunoblotting

Samples from *in vitro* binding assays were separated by SDS PAGE on GenScript 4-20% Bis-Tris gel and transferred to PVDF membrane (0.45 μm) for 1h 20 min at 35 V using Invitrogen mini blot module. For protein detection, membranes were incubated with primary mouse monoclonal anti-GFP (Roche) or goat polyclonal anti-GST antibody (GE Healthcare) and HRP-conjugated secondary antibodies (Dianova) followed by treatment with enhanced chemiluminescence substrate kit (ECL, GE Healthcare). Images were acquired with LAS4000 (GE ImageQuant) or ImageQuant 600 (Cytiva).

##### AlphaScreen proximity assay

Ala-peptide binding to Pirh2 or KLHDC10 was evaluated using the AlphaScreen GST detection Kit (PerkinElmer) proximity-based assay. The two assay components, streptavidin-coated donor beads that associate with biotin-labeled peptide and anti-GST acceptor beads that associate with GST-fused protein, are brought in proximity depending on peptide-protein interaction. Upon excitation of donor beads, energy transfer to acceptor beads takes place that generates a luminescence signal. Experiments were performed in ProxiPlate 384-well microplates (PerkinElmer).

Competition experiments to determine IC_50_ were performed by titrating unlabeled peptides. All required dilutions of reaction components were made in assay buffer (20 mM Tris pH 7.5 at room temperature, 100 mM NaCl, 0.1% Tween-20, 0.05% BSA). 10 nM biotin-labeled Ala-tail peptide (MDELYKAAAAAA) (Biosyntan) and 1 nM GST-Pirh2 or GST-KLHDC10 were pre-incubated with varying concentrations of competing unlabeled Ala-tail variant peptide (GenScript) for 40 min at RT. Following this incubation, pre-mixed donor and acceptor beads (10 μg/ml each) were added to the reaction mixture, in a total 20 μl, and further incubated for 40 min at RT. AlphaScreen signal was measured using PHERAstar microplate reader (BMG Labtech) at RT. Each reaction was performed in triplicates and IC_50_ was determined by non-linear curve fitting of dose response curve in GraphPad Prism 8.

##### Structure predictions using AlphaFold2

Peptide bound complexes of Pirh2 and KLHDC10 were predicted using AlphaFold2.^36, 37^ The experiments were run through ColabFold^52^ assessed through UCSF ChimeraX 1.4 and 1.5.^53^ For one experiment, sequences of Pirh2_1-195_ or KLHDC10_β1-88-442_ and Ala-tail (MDELYKAAAAAA) were input separately. In a second experiment, the Ala-tail peptide sequence was fused to the C-terminus of the protein following a Gly_50_ linker. For KLHDC10, complexes were also predicted using a shorter Ala-tail peptide (LYKAAAAAA). Default settings without any templates from PDB database were used to run the predictions. For every run, the top 5 predictions were obtained as output from the server. Models were analyzed using UCSF Chimera.^54^

##### Sequence analysis

The sequences of the mouse Pirh2 and human KLHDC10 were obtained from the National Center for Biotechnology Information (NCBI) Genbank database^55^ and dissected into constituent domains using known structures and reverse searches against sequence profiles. Sequence similarity searches were performed using the PSI-BLAST program^56^ against the NCBI non-redundant (nr) database or the same database clustered down to 50% sequence identity using the MMseqs program^57^ with a profile-inclusion threshold set at an e-value of 0.01. Profile-profile searches were performed with the HHpred program.^58^ Multiple sequence alignments (MSAs) were constructed using the FAMSA^59^ and MAFFT programs.^60^ Multiple sequence alignments (MSA) of Pirh2 and KLHDC10 homologs were generated using Clustal Omega^61^ and displayed using ESPript 3.0.^62^ Conserved residues were mapped onto the AlphaFold predicted structures of full-length human Pirh2 and KLHDC10 using ConSurf webserver (https://consurf.tau.ac.il/consurf_index.php).^63^

##### Structure analysis

The JPred program was used to predict secondary structures using MSAs (see above). PDB coordinates of structures were retrieved from the Protein Data Bank and rendered, compared, and superimposed using the Mol* program.^64^ Structural models were generated using the RoseTTAfold^65^ and Alphafold2 programs.^36^ Multiple alignments of related sequences (>30% similarity) were used to initiate HHpred searches for the step of identifying templates to be used by the neural networks deployed by these programs.

##### Comparative genomics and phylogenetic analysis

Phylogenetic analysis was performed using the maximum-likelihood method with the WAG or LG models with the IQTree^66^ and FastTree program,^67^ with 8-20 rate categories and 1 invariant category for the sites. The FigTree program (http://tree.bio.ed.ac.uk/software/figtree/) was used to render phylogenetic trees. Gene neighborhoods were extracted through custom PERL scripts from genomes retrieved from the NCBI Genome database. These were then clustered using MMSEQS adjusting the length of aligned regions and bit-score density threshold empirically and filtered using neighborhood distance cutoffs and phyletic patterns to identify conserved gene neighborhoods.

#### Quantification and statistical analyses

Data were plotted using GraphPad Prism 8. Error bars represent standard deviation.

#### Supplemental information

Figures S1-S10

