## Supplementary material for "Mechanism and evolutionary origins of Alanine-tail C-degron recognition by E3 ligases Pirh2 and CRL2-KLHDC10": Figures S1-S10

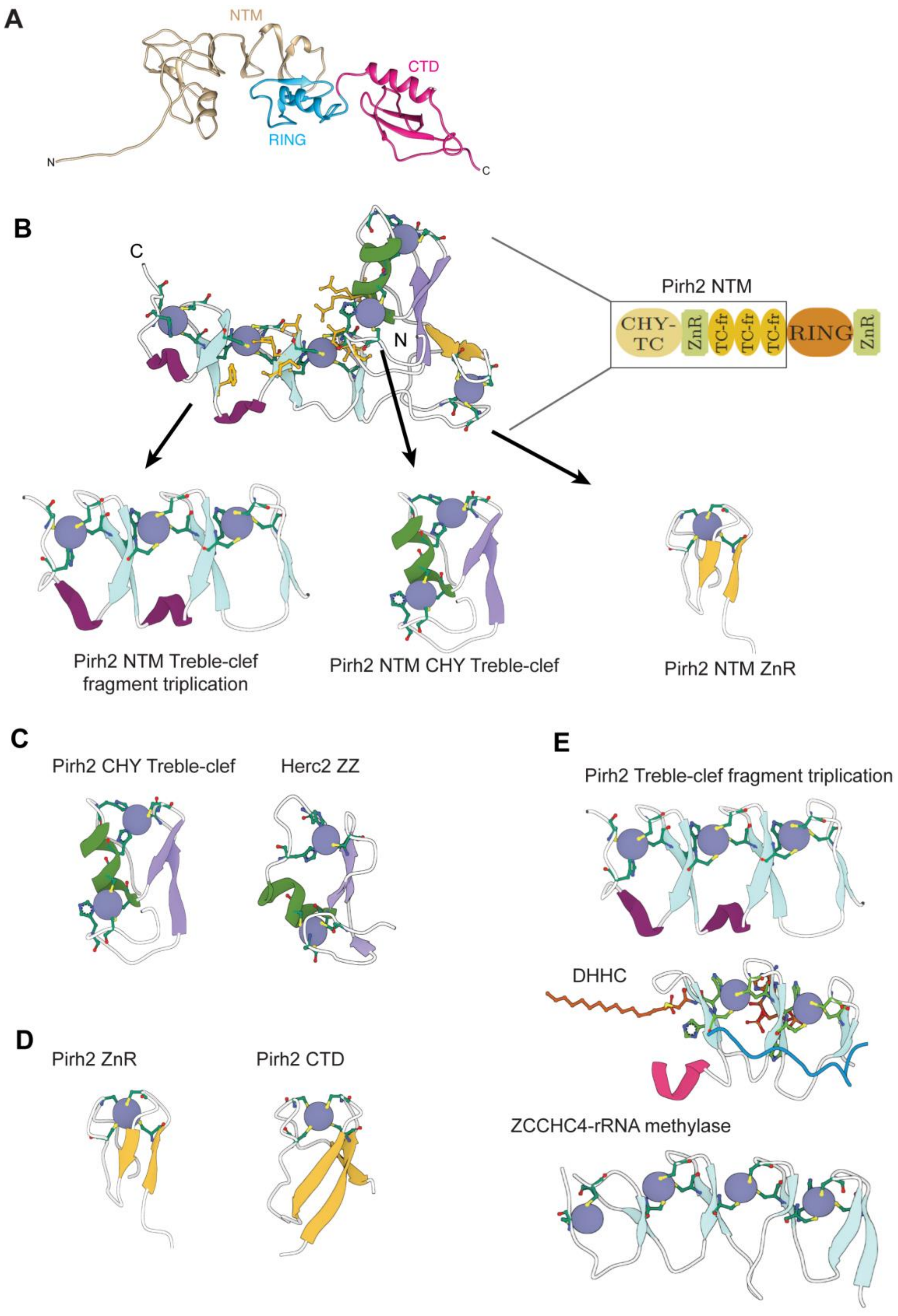


**Figure S1. Human Pirh2 structure.**
(A) Structure of full-length Pirh2 (UniProt: Q96PM5) predicted by AlphaFold. Regions of the protein are colored according to Figure 1A. The NTM consists of a short unstructured region of 17 amino acids

at the very N-terminus followed by zinc binding domains (see ‘B’). The CTD consists of a classic Zn-ribbon (ZnR) domain (see ‘B’).

(B) N-terminal module (NTM) of Pirh2 showing its composite structure and constituent domains, from N-terminus to C-terminus: CHY Treble-clef (CHY-TC), classic Zn-ribbon (ZnR), and Treble-clef fragment (TC-fr) triplication. Note that the Pirh2 NTM structure is shown in opposite orientation compared to that in panel ‘A’.

(C) The N-terminal most CHY Treble-clef domain compared to a homologous histone H3.1 binding ZZ domain from the HECT-type E3 ligase HERC2 (PDB: 6WW4).

(D) The ZnR domain compared to the homologous domain found in the CTD of Pirh2 (PDB:2K2D).

(E) The three C-terminal repeats are fragmentary versions of the Treble-clef comprised of just the C-terminal part of this domain. It is compared to homologous Treble-clef fragments that are found in the catalytic domain of the DHHC palmitoyltransferases (PDB: 6BML) and in the RNA-binding domain of the 28S rRNA adenine methylase ZCCHC4 (PDB: 6UCA).

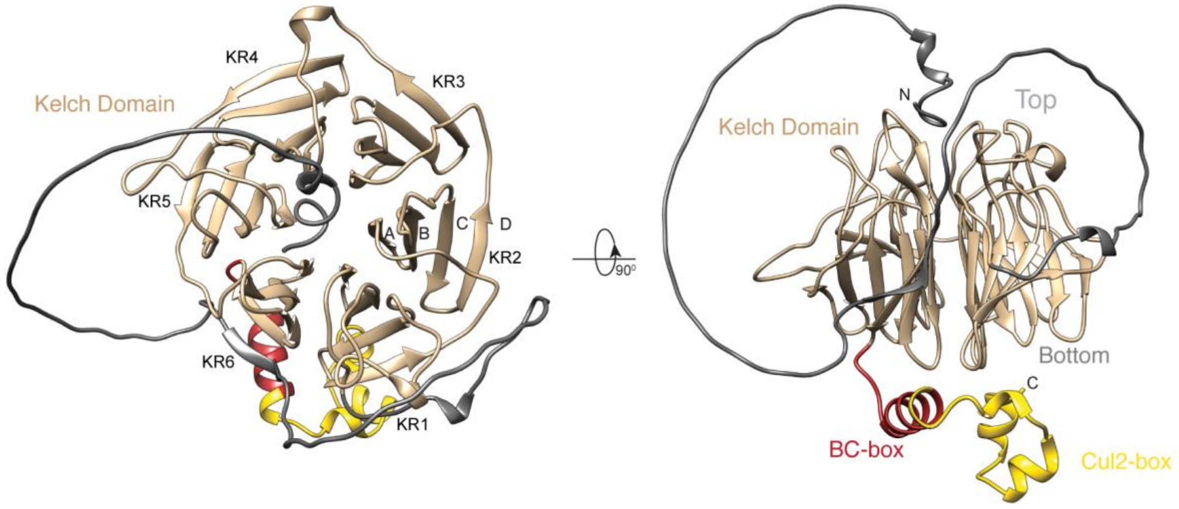


**Figure S2. Human KLHDC10 structure.**
Views from top (*left*) and side (*right*) of the AlphaFold predicted model for full-length human KLHDC10 (UniProt: Q6PID8). Domains are colored according to Figure 1D. The β-propeller domain in KLHDC10 is formed by 6 Kelch repeats, each folding as a propeller blade. In turn, each blade is formed by four β-strands, labelled as A-D as indicated for Kelch Repeat 2 (KR2).

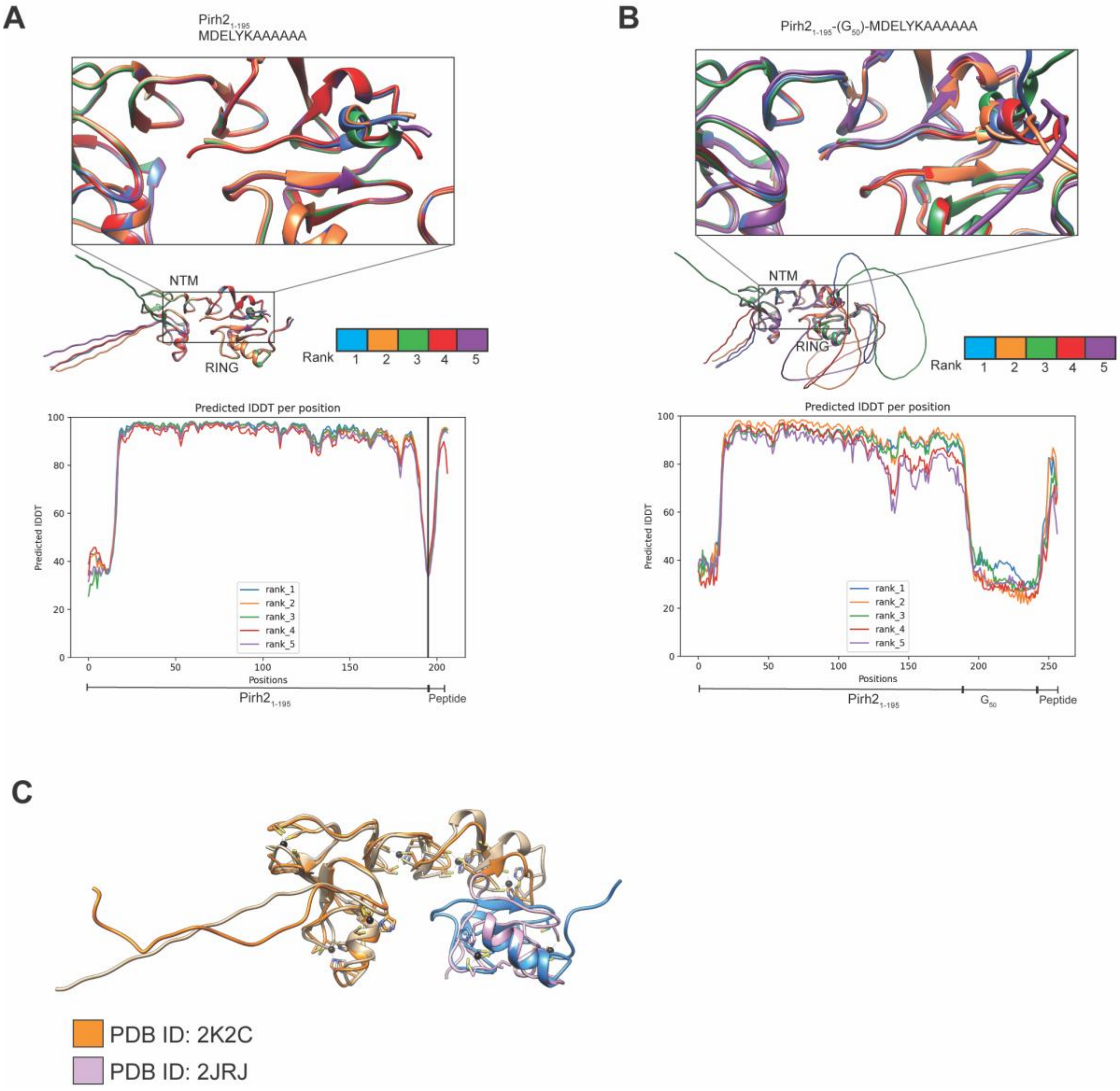


**Figure S3. AlphaFold2-predicted complexes of Pirh2$_{1-195}$ and an Ala-tail peptide.**
(A) Superposition of the top five models obtained when Pirh2$_{1-195}$ and Ala-tail peptide sequences were separately submitted (i.e., in *trans*) for complex prediction. Each complex is colored according to the key presented, with rank 1 being the top scoring model. The binding site is also shown in close-up view. *Below*, graph of pLDDT score per residue of each predicted model.
(B) As in panel A, but superposition of the top five models obtained when Pirh2$_{1-195}$ and Ala-tail peptide sequences were submitted as a fusion protein (i.e., in *cis*) for complex prediction.
(C) Superposition of the top scoring Pirh2$_{1-195}$ AlphaFold prediction from panel A with individual NMR structures of the Pirh2 NTM (orange) and RING domain (purple), to indicate Zinc binding sites (zinc ions represented as back circles).

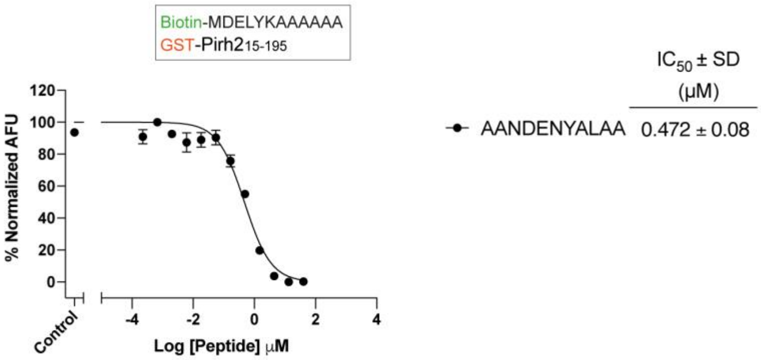


**Figure S4. Pirh2 directly binds to an SsrA peptide.**
As in Figure 1C, except that the competing peptide sequence was AANDENYALAA, corresponding to the *B. subtilis* SsrA tag.

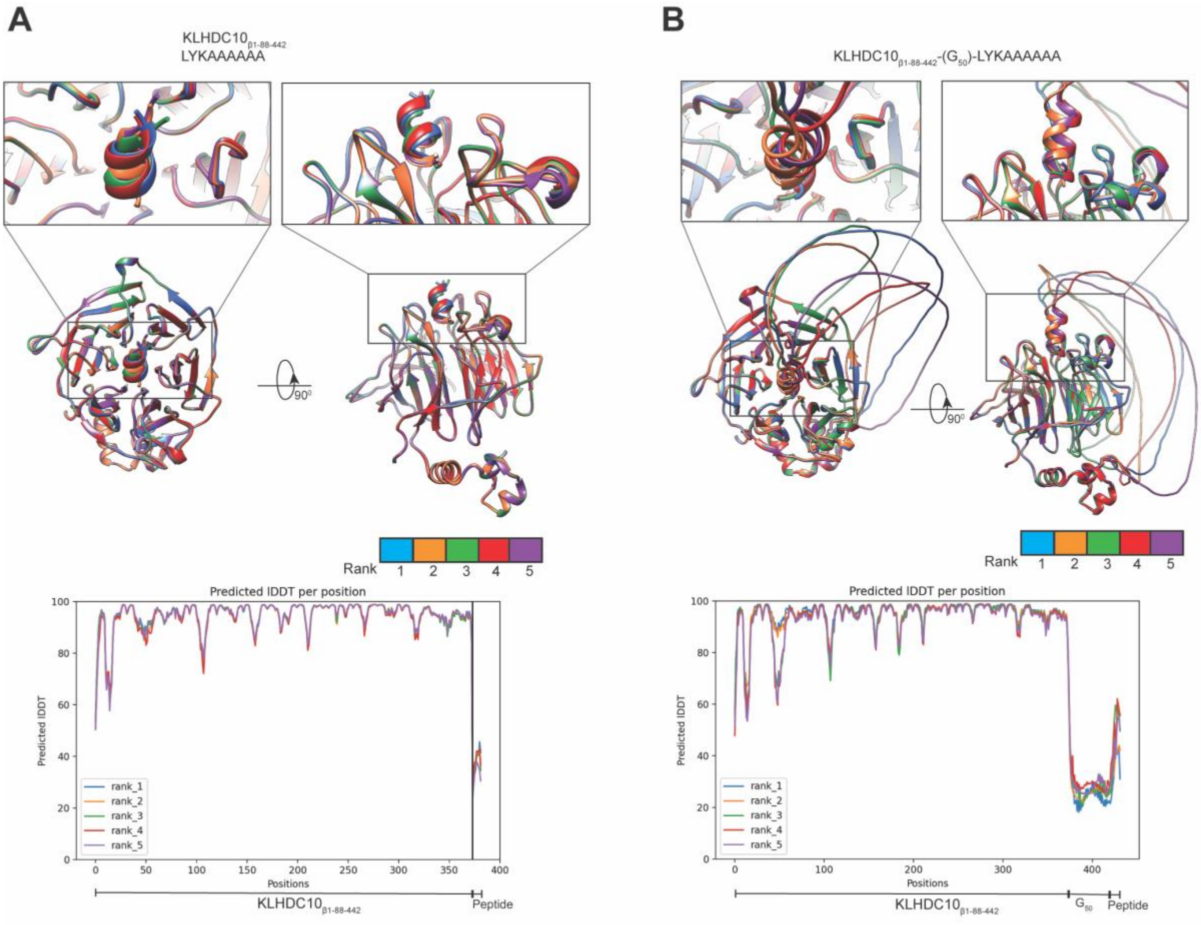


**Figure S5. AlphaFold2 predicted complexes for KLHDC10$_{\beta 1\text{-}88\text{-}442}$ and an Ala-tail peptide.** KLHDC10 views as in Figure S2B. (A) Superposition of the top five models obtained when KLHDC10$_{\beta 1\text{-}88\text{-}442}$ and a shorter Ala-tail peptide (LYKAAAAAA) sequence were separately submitted for complex prediction. Each complex is colored according to the key presented, with rank 1 being the top scoring model. The binding site is also shown in close-up view. *Below*, graph of pLDDT score per residue of each predicted model.

(B) As in A, but superposition of the top five models obtained when KLHDC10$_{\beta 1\text{-}88\text{-}442}$ and a shorter Ala-tail peptide sequences were submitted as a fusion protein for complex prediction.

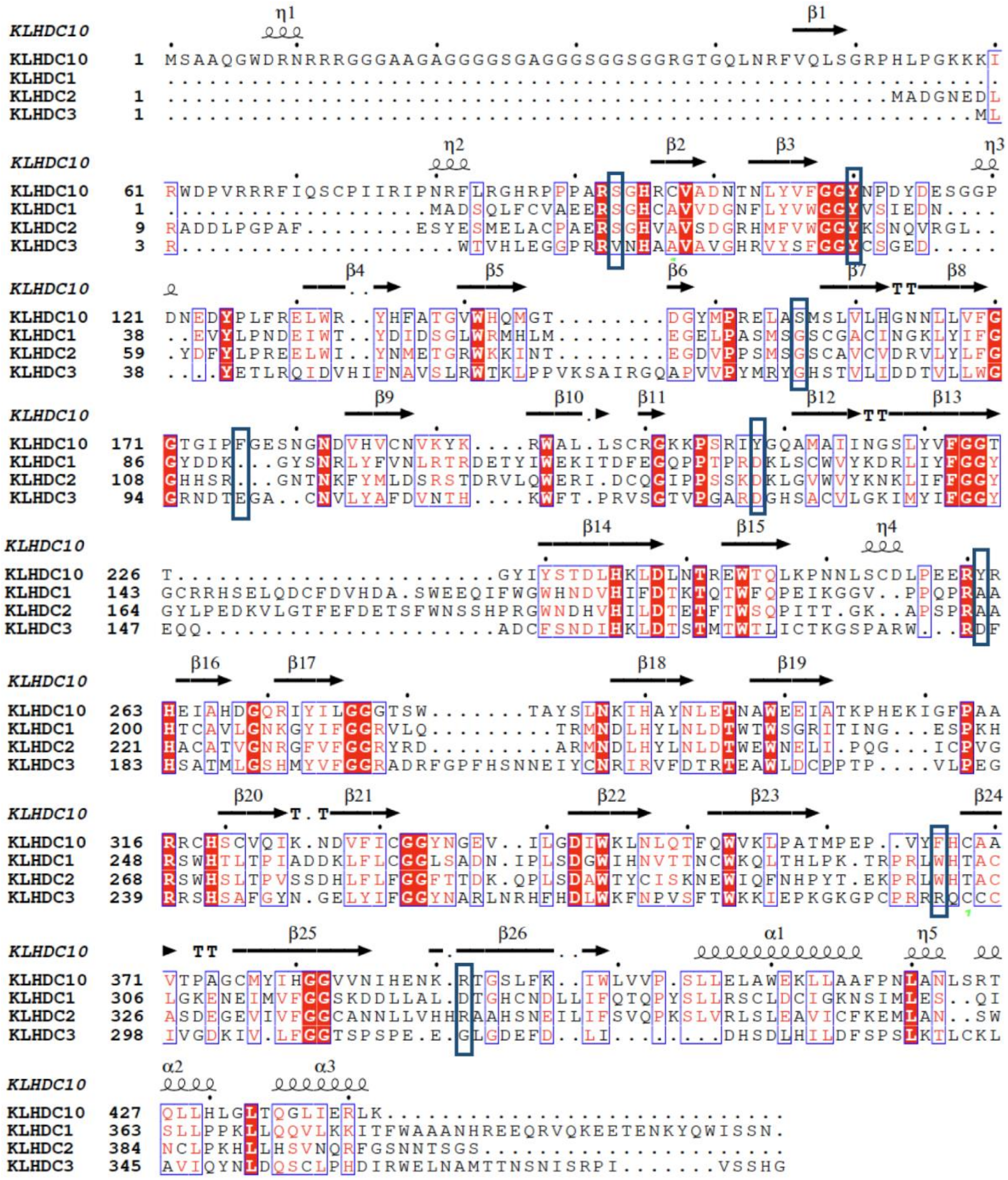

**Figure S6. Sequence comparison of human KLHDC proteins.**
Blue boxes highlight some residues in KLHDC10 implicated in Ala-tail recognition and residues at the corresponding positions in other KLHDC proteins.

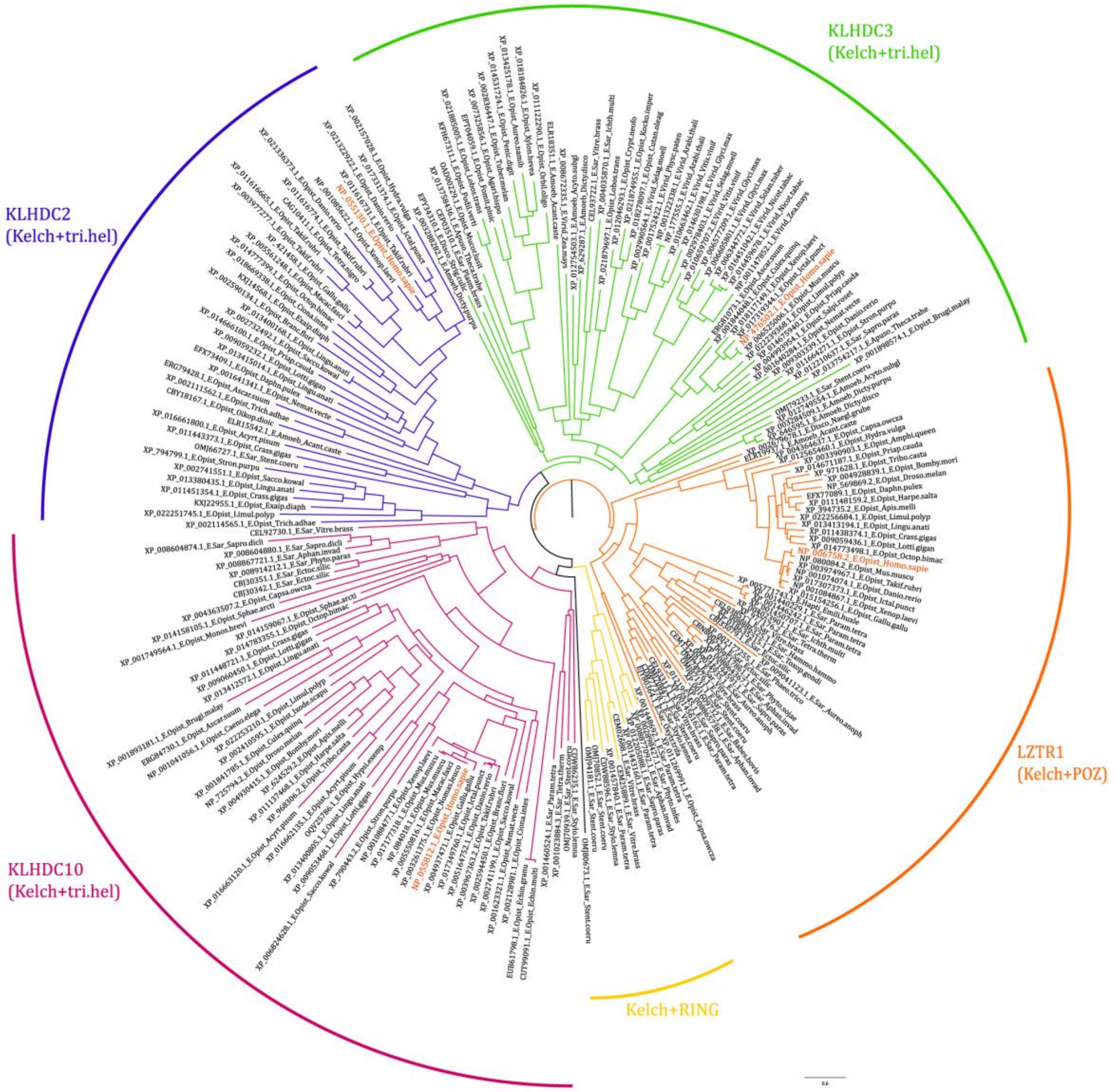


**Figure S7. A tree of E3 Kelch domain β-propellers**.
The clades are colored differently and labeled according to their human representatives (KLHDC10, KLHDC2, KLHDC3, LZTR1) or domain architecture (the Kelch+RING proteins of certain microbial eukaryotes). The archetypal human proteins of each clade are shown in bold and red in the tree. All the labeled clades are supported by bootstrap support of > 90%.

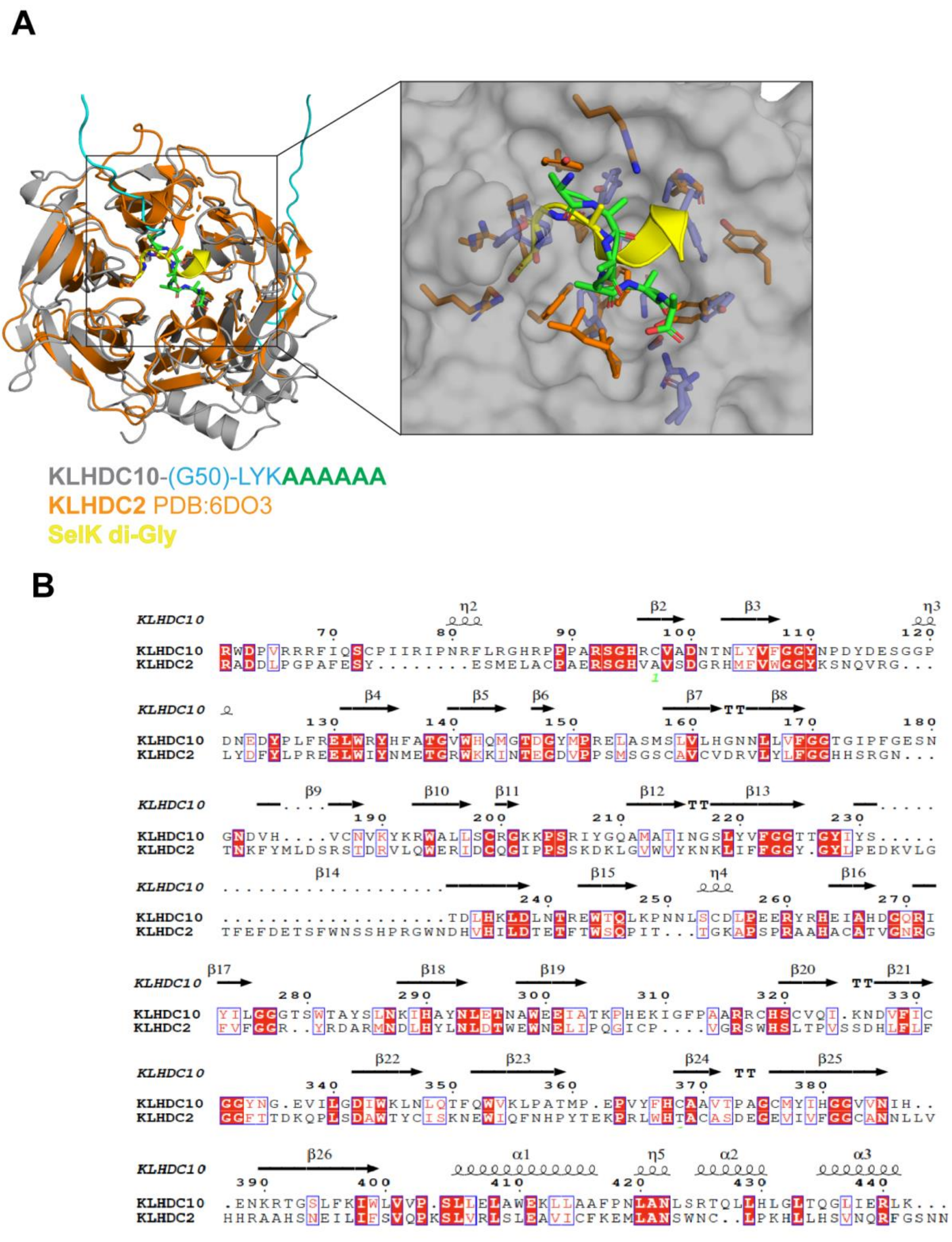


**Figure S8. KLHDC10 and KLHDC2 show discrete C-degron recognition modes.**
(A) Superposition of AlphaFold predicted Ala-tail degron recognition model of KLHDC10 and the co-crystal structure of KLHDC2 with SelK di-Gly C-degron. In the inset, Ala-tail peptide (in green) and SelK di-Gly degron (in yellow); surface of KLHDC10 represented in grey.
(B) Sequence alignment of human KLHDC10 and KLHDC2.

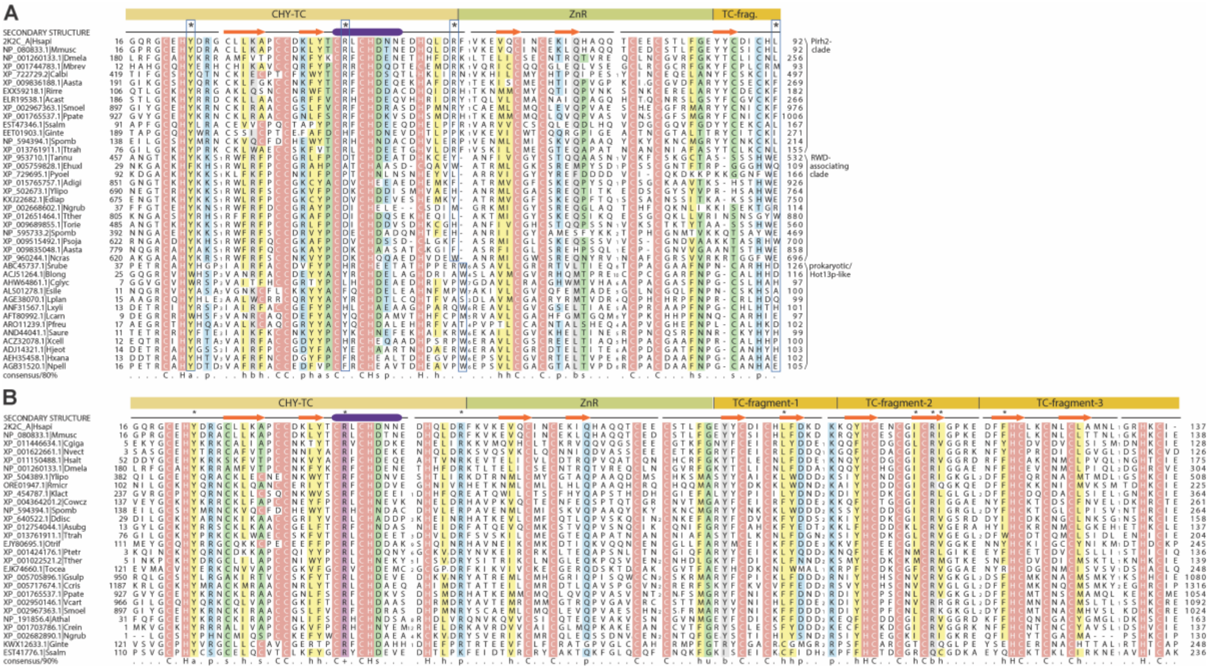

**Figure S9**.
Multiple sequence alignments of the (A) core CHY-TC (CHY-Treble Clef) and ZnR module found in the NTM and (B) the same module in the Pirh2-like proteins with their adjacent TC (Treble Clef)-fragment repeats. Individual domain boundaries are marked at top of the alignment. Monophyletic clades of proteins are labeled to the right in (A). Conserved amino acid residues are colored according to their side chain chemistry or geometry, with metal-chelating residues shaded in red and colored in white. The consensus sequence was calculated based on a larger set of 294 representatives (A) and 160 representatives (B). The residue classes are defined thus: a: aromatic; b: big; h: hydrophobic; s: small; p: polar; u: tiny; +: basic. Positions identified as contacting residues in this study are marked above the alignment with asterisks. These positions in A are further boxed in blue for emphasis. Sequences are labeled by NCBI accession number, with species name abbreviation as follows: (A) Aastac: *Aphanomyces astaci*; Acast: *Acanthamoeba castellanii*; Adigi: *Acropora digitifera*; Blong: *Bifidobacterium longum*; Calbi: *Candida albicans*; Cglyc: Corynebacterium glyciniphilum; Dmela: *Drosophila melanogaster*; Ediap: *Exaiptasia diaphana*; Ehuxl: *Emiliania huxleyi*; Esile: *Enterococcus silesiacus*; Ginte: *Giardia intestinalis*; Hjeot: *Halalkalicoccus jeotgali*; Hxana: *Halopiger xanaduensis*; Lcarn: *Leuconostoc carnosum*; Lplan: *Lactiplantibacillus plantarum*; Lxyli: *Leifsonia xyli*; Mbrev: *Monosiga brevicollis*; Mmusc: *Mus musculus*; Ncras: *Neurospora crassa*; Ngrub: *Naegleria gruberi*; Npell: *Natrinema pellirubrum*; Pfreu: *Propionibacterium freudenreichii*; Ppate: *Physcomitrella patens*; Psoja: *Phytophthora sojae*; Pyoel: *Plasmodium yoelii yoelii*; Rirre: *Rhizophagus irregularis*; Saure: *Staphylococcus aureus*; Smoel: *Selaginella moellendorffii*; Smuta: *Streptococcus mutans*; Spomb: *Schizosaccharomyces pombe*; Srube: *Salinibacter ruber*; Ssalm: *Spironucleus salmonicida*; Tannu: *Theileria annulata*; Torie: *Theileria orientalis*; Tther: *Tetrahymena thermophila*; Ttrah: *Thecamonas trahens*; Xcell: *Xylanimonas cellulosilytica*; Ylipo: *Yarrowia lipolytica*. (B) Asubg: *Acytostelium subglobosum*; Athal: *Arabidopsis thaliana*; Ccris: *Chondrus crispus*; Cgiga: *Crassostrea gigas*; Cowc: *Capsaspora owczarzaki*; Crei: *Chlamydomonas reinhardtii*; Ddisc: *Dictyostelium discoideum*; Gsulp: *Galdieria sulphuraria*; Hsalt: *Harpegnathos saltator*; Hsapi: *Homo sapiens*; Klact: *Kluyveromyces lactis*; Nvect: *Nematostella vectensis*; Otrif: *Oxytricha trifallax*; Ptetr: *Paramecium tetraurelia*; Rmicr: *Rhizopus microsporus*; Tocea: *Thalassiosira oceanica*; Vcart: *Volvox carteri*.

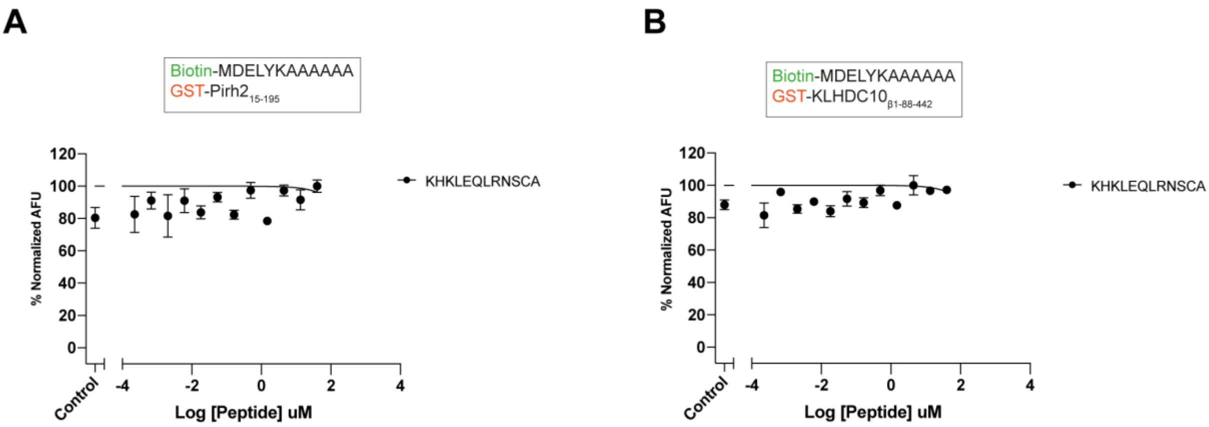


**Figure S10. Pirh2 and KLHDC10 does not bind to C-end peptide of Myc protein.**
As in Figure 1C, except that the competing peptide sequence was KHKLEQLRNSCA, corresponding to the C-terminal sequence from human protein c-Myc tested for its binding to A) Pirh2 and B) KLHDC10.
